# Oncogenic and Circadian Effects of Small Molecules Directly and Indirectly Targeting the Core Circadian Clock

**DOI:** 10.1101/645861

**Authors:** Hui-Hsien Lin, Kelly L. Robertson, Heather A. Bisbee, Michelle E. Farkas

**Affiliations:** Department of Chemistry, University of Massachusetts, Amherst, MA; Department of Biochemistry and Molecular Biology, University of Massachusetts, Amherst, MA; Molecular and Cellular Biology Graduate Program, University of Massachusetts, Amherst, MA

**Keywords:** Circadian rhythms, small molecules, clock periods, U2OS cells, cell proliferation, cell migration, tumor growth

## Abstract

Circadian rhythms are essential for controlling the cell cycle, cellular proliferation, and apoptosis, and hence, are tightly linked to cell fate. Disruption of circadian rhythms has been shown to trigger various pathological developments, including cancer. Several recent studies have used a variety of small molecules to affect circadian oscillations, however, their concomitant cellular effects were not assessed. Here, we use five molecules, grouped into direct versus indirect effectors of the circadian clock, to modulate periods in a human osteosarcoma cell line (U2OS), and determined their influences on cellular behaviors, including motility and colony formation. Luciferase reporters, whose expression were driven via *Bmal1*- and *Per2*-promoters (positive and negative protein components of the core clock), were used to facilitate the visualization and quantitative analysis of circadian oscillations. We show that all molecules significantly increase or decrease the circadian periods of *Bmal1* and *Per2* in a dose-dependent manner, but period length does not correlate with the extent of cell migration or proliferation. We observed that only molecules that affected circadian oscillations to a greater extent showed significant influence on cell functions (e.g. motility and colony formation). Because it is important to consider the likelihood of biological effects resulting from non-circadian targets, we also provide a thorough discussion of potential modes of action. Future studies should employ additional compounds that directly target circadian proteins and/or have different circadian effects, and evaluation in other cancer models to determine whether results obtained here remain consistent.

**For Table of Contents Only:** 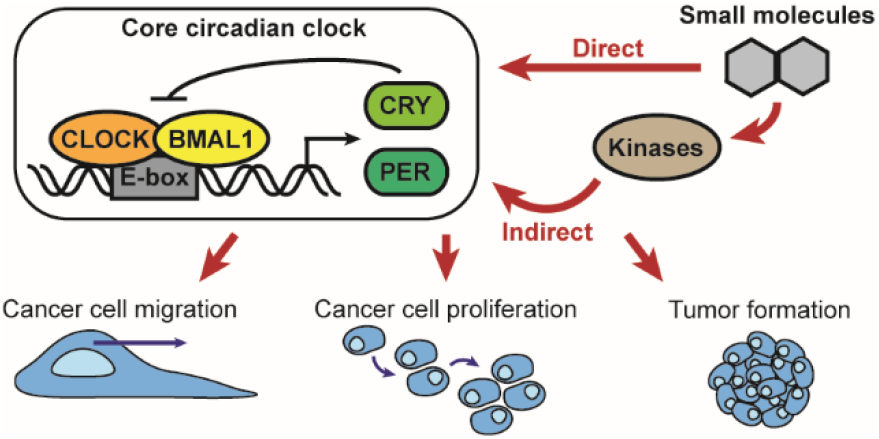

## Introduction

Circadian rhythms are endogenous time-keeping systems that cycle with an approximate period of 24 h.^1^ These autonomous clocks help organisms adapt to environmental changes, such as day-night cycles^2^ or the changing of seasons.^3^ Light and temperature inputs regulate circadian rhythms through the hypothalamic suprachiasmatic nucleus (SCN), which is the master clock regulator for the entire organism.^4^ Circadian clocks coordinate daily changes to control many physiological, behavioral, and metabolic functions, including the sleep-wake cycle,^5^ body temperature,^6^ blood pressure,^7^ food intake,^8^ and humoral secretion.^9^ Disruption of circadian rhythms via night shift-work,^10^ exposure to light at night,^11^ or chronic jet lag^12^ has been shown to elicit a number of pathological developments, including cancer,^13^ metabolic disorders,^14^ and cardiovascular diseases.^15^ While accumulating evidence indicates that circadian disruptions lead to malignant transformations in cells and organs, the molecular relationship between the two is not well-understood.

The circadian system in mammals is hierarchically organized into two major components: a central pace-maker SCN and peripheral oscillators within cells.^16^ When the retina receives photons from the environment, it signals the SCN to synchronize peripheral clocks via neural and humoral pathways.^17^ At the cellular level, molecular clocks are regulated by transcriptional/translational feedback loops (TTFLs) integrated in clock genes. Core clock proteins aryl hydrocarbon receptor nuclear translocator-like protein 1 (ARNTL/BMAL1) and circadian locomotor output cycles kaput (CLOCK) heterodimerize in the morning and bind to the E-box promoter, which initiates the expression of *Period* (*Per*) and *Cryptochrome* (*Cry*). PER and CRY accumulate in the evening and form a heterodimer, which translocates back into the nucleus and inhibits *Bmal1* and *Clock* activity.^18^ As a result, the core circadian machinery oscillates with a period of approximately 24 h. In cell culture, intrinsic and self-sustained circadian clocks are persistent even in the absence of external time cues.^19^ To synchronize clocks in cultured cells, treatments of high concentration serum^20^ or chemical reagents (e.g. dexamethasone^21^ or forskolin^22^) are frequently used.

Circadian rhythmicity is tightly associated with post-translational modifications of clock proteins.^23^ Phosphorylation of most clock proteins occurs in a rhythmic manner; thus, alteration of clock protein phosphorylation can result in changes to circadian periods.^24^ Genetic manipulations of the post-translational regulators of clock proteins have been shown to affect circadian functions and periodicity.^24–25^ However, conventional genetic approaches also result in fatality, pleiotropy, and functional redundancy. Alternatively, chemical modulation of these regulators via small molecules can reversibly manipulate clock functions in time- and dose-dependent manners.

Recently, the identification of small molecule modulators of circadian rhythms has garnered much interest,^26^ but few directly interact with clock proteins, including CRYs,^26b,27^ REV-ERBs,^28^ and RORs.^29^ Most of the others target kinases (e.g. CK2, GSK3β, and AMPK)^30^ that regulate clocks via post-translational modifications.^31^ These molecules have promoted the study of post-translational mechanisms underlying the circadian system and been used to discover new clock-regulatory pathways.^23a,32^ Some circadian-modulating molecules have shown therapeutic potential as well. It has been recently reported by Sulli et al. that SR9009 and SR9011, two agonists of REV-ERBs (secondary clock components), have potent anti-cancer effects in various cancer cell lines, and no toxicity in normal cells or tissues.^33^ Oshima et al. discovered GO289, which regulates circadian rhythms via inhibition of CK2, finding that it suppressed cancer cell proliferation in a cell type-dependent manner.^30a^ But, among work published, the biological and circadian effects of most small molecules, including those studied here, have not been addressed concurrently, nor have they been studied in similar systems using the same modes of analysis. Hence, we present a systematic assessment, whereby we have thoroughly evaluated a small panel of these molecules, also addressing whether concentration ranges resulting in circadian effects yield cellular changes and vice-versa.

In this study, we used five small molecules to affect circadian period (the time interval for completing one oscillation cycle), and assessed concomitant changes to the oncogenic traits of cancer cells. Among the selected compounds, two, KL001^27^ and PF-670462,^34^ result in direct effects on circadian proteins and the other three, SP600125,^35^ Chir99021, and Etoposide,^36^ target entities that regulate core circadian proteins in a post-translational manner (Fig. 1). A human bone osteosarcoma epithelial cell line, U2OS, was separately transfected with *Bmal1:luc* and *Per2:luc* reporters to facilitate high resolution tracking of circadian rhythms, and used throughout this work. We confirmed that the five small molecules all either increase or decrease the period of *Bmal1* and *Per2* in these cells, to varying degrees. However, we found that conditions resulting in circadian effects did not necessarily result in changes to oncogenic features. This leads us to posit that circadian periods may not be correlated with cell motility or growth. While KL001, PF-670462, and SP600125 all increased period length, KL001 slightly promoted cell migration while the other two inhibited it. This contradiction was also observed between Chir99021 and Etoposide treatments, which both decreased periods, but while Chir99021 suppressed cell migration, Etoposide had no effect. Our data suggest that there may not be a direct connection between circadian rhythms and cancer cell behavior, and the circadian period length may not be a key regulator of cell motility and growth. To confirm whether this is the case, and in further studies using small molecules to affect and study circadian rhythms, molecules directly targeting additional core clock components should be identified and employed. These may be combined with genetic approaches to uncover the molecular details connecting circadian disruptions with cancer development.

**Figure 1.**
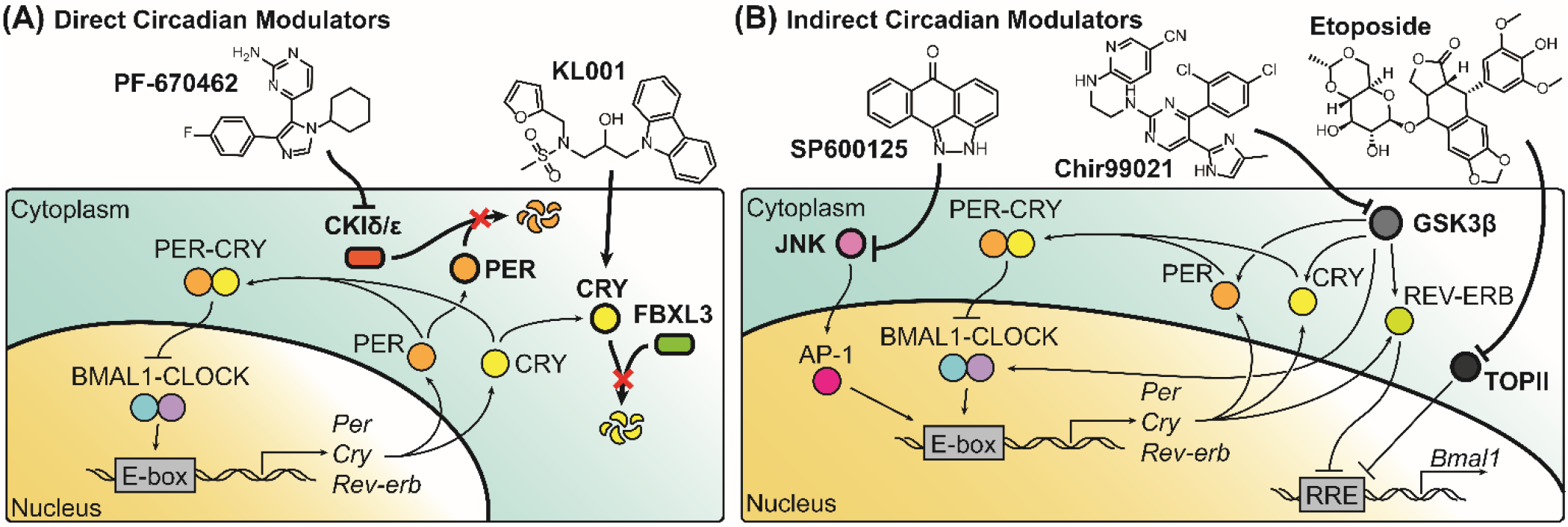
Structures and targets of direct circadian modulators (**A)** and indirect circadian modulators (**B)** used in these studies. KL001 directly binds to CRY, preventing FBXL3-mediated degradation, and PF-670462 binds CKIδ/ε resulting in PER stabilization. SP600125, Chir99021, and Etoposide, indirectly influence circadian rhythms by binding to kinases that are involved in circadian pathways.

## Materials and Methods

### Cell Culture

U2OS cell line was obtained from Prof. Patricia Wadsworth (Biology, UMass Amherst). Cells were maintained in DMEM (Gibco 11960-044), with 10% Fetal Bovine Serum (FBS; Corning), 10% L-Glutamine (Gibco), 1% penicillin-streptomycin (Gibco), 1% Non-Essential Amino Acids (HyClone), and 1% Sodium Pyruvate (Gibco). HEK293T cell line was obtained from Prof. D. Joseph Jerry (Veterinary and Animal Sciences, UMass Amherst). Cells were maintained in DMEM/F12 (Gibco 11320-033), with 10% FBS, 1% penicillin-streptomycin (Gibco), and 0.015 mg/mL gentamicin (Gibco). All cells were incubated at 37 °C under 5% CO_2_, except where otherwise noted.

### Plasmid and Recombinant DNA

The *Bmal1* promoter driven luciferase reporter construct (pABpuro-BluF, *Bmal1:luc*) was obtained from Addgene (plasmid #46824, deposited by Dr. Steven Brown).^37^ Our generation of the *Per2* promoter driven luciferase (*Per2:luc*) reporter construct was described previously.^38^

### Lentiviral Transductions

3 x 10^6^ HEK293T cells were seeded in 60 mm culture dishes and transiently transfected with 3 μg psPAX packaging plasmid, 2 μg pMD2G envelope plasmid (both from Prof. D. Joseph Jerry, Veterinary and Animal Sciences, UMass Amherst), and 3 µg *Bmal1:luc* or *Per2:luc* reporter constructs using Lipofectamine3000 (ThermoFisher Scientific), according to the manufacturer’s instructions. Lentiviral particles were harvested from supernatant 48 h after DNA-lipid complexes were added to cells; this virus-containing supernatant was passed through a 45 µm filter. 9 mL lentivirus-containing supernatant was mixed with 9 mL DMEM culture medium containing 10 µg/mL polybrene (Sigma). U2OS cells were seeded in T25 culture flasks at 2 x 10^5^ cells/mL and incubated under conditions above until 70-80% confluence was reached. Culture medium was removed and 6 mL of lentiviral-containing media was added to each flask. After two days of infection, the medium was replaced with selection medium (DMEM with all growth supplements plus 4 µg/mL puromycin), in which cells were incubated for 3-6 weeks for selection.

### Bioluminescence Recording

Cells were seeded in 35 mm culture dishes with 2 mL of 2 x 10^5^ cells/mL and incubated to reach 100% confluence. After 24 h, culture media was replaced with bioluminescence recording media, (powdered DMEM (Sigma-Aldrich) with 4 mM sodium bicarbonate (Gibco), 5% FBS (Corning), 1% HEPES (HyClone), 0.25% penicillin-streptomycin (Gibco), 53.5 mM d-Luciferin (ThermoFisher)). 100 nM dexamethasone was added to recording media for synchronization. Dishes were sealed with 40 mm sterile cover glass using silicon vacuum grease and subjected to monitoring using a LumiCycle 32 System (Actimetrics) at 36.5 °C for 5-7 days. Raw traces were detrended using the 24-h moving average method via LumiCycle Analysis v.2.56 software. All period analyses were performed via simultaneous Levenberg-Marquardt least-squares parameter optimization with damped sinusoidal waveform using the same software.

### Small Molecule Cell Treatments

SP600125 (Sigma-Aldrich), KL001 (Tocris), PF-670462 (Sigma-Aldrich), Chir99021 (Tocris), and Etoposide (Santa Cruz Biotech), were prepared in 100% cell culture grade dimethyl-sulfoxide (DMSO; Sigma-Aldrich) at a concentration of 50 mM and stored at −20 °C in single use aliquots. When dosing cells, each was diluted in culture media (or recording media for bioluminescence recording) to a final DMSO concentration of 0.2%. The solution was mixed well and added to the seeded cells.

### Wound Healing Assay

Cells were seeded in 24-well plates at 2 x 10^5^ cells/mL and incubated for 24-48 h. After cells reached 100% confluence, wounds were generated using a 1 mL micropipette tip (Denville). Culture media was removed, cells were washed with 500 µL PBS (Gibco), and 500 µL of new culture media containing indicated molecules was added into each well. Images were taken immediately following for the first time point (T = 0) and then every 6 h for 24 h via Biotek Cytation 3 cell imaging multi-mode reader. Wound closure rates were determined by normalizing wound area at each time point to T = 0 via ImageJ.

### Colony Formation Assay

Cells were suspended in agar and incubated until colonies formed, as previously described.^39^ Briefly, 3% 2-Hydroxyethyl Agarose (Sigma) was prepared and stored in a water bath at 45 °C. Cell culture media at 37 °C was used to dilute agarose to 0.6%, and 2 mL was added to each well of a 6-well plate (Nuclon), which was incubated at 4 °C until agar solidified (first, bottom layer). DMSO solutions of molecules were added to 1 mL of 4 x 10^4^ cells/mL suspended U2OS cells in warm culture media, followed by addition of 0.6% agarose solution in a 1:1 ratio. 1 mL drug-cell-agarose solution was dispensed per well and incubated at 4 °C for 30 min until the agar layer solidified (second, drug-cell containing layer). Then plates were incubated at 37 °C under 5% CO_2_. A 1 mL feeder layer of 0.3% agarose in culture media containing the small molecules at designated concentrations was added to each well once every 7 days for 4 weeks. When imaging colonies, 1 mL of 0.005% Crystal Violet dye (Fisher) in 20% methanol (Fischer Scientific) was added to wells and incubated on a shaker at rt for 1 h. Wells were then washed 4-6 times with 2 mL ddH_2_O. Eight images were taken per well using a Biotek Citation 3 multi-mode cell imaging reader. Each condition was performed in three biological replicates. Images were stitched using Adobe Illustrator CS6 and colony numbers/sizes were analyzed using ImageJ.

## Results and Discussion

### Both direct and indirect circadian modulators alter periods in a dose-dependent manner

To evaluate the changes to circadian period caused by the five small molecules used, real-time luminometry assays were performed using human U2OS osteosarcoma cells harboring *Bmal1:luc* or *Per2:luc* reporters (**Fig. S1**). Among the molecules, KL001^27^ and PF-670462^34^ are categorized as direct circadian modulators, because their targets, CRY and CKIδ/ε, are essential elements for completing endogenous circadian TTFLs. SP600125,^35, 40^ Chir99021, and Etoposide^36^ are classified as indirect circadian modulators, because their targets, JNK, GSK3β, and TOPII, exert down-stream effects on endogenous circadian TTFLs.

KL001 directly binds to CRY, inhibiting F-box and leucine-rich repeat protein 3 (FBXL3)-mediated CRY degradation.^27^ Treatment with KL001 was observed to both increase period and dampen *Bmal1:luc* amplitude in a dose-dependent manner (**Figs. 2A, 2B, S2,** and **Table S1)**, within the IC50 range (0.82 to 14 μM).^27^ While perturbation of the FBXL3-CRY regulatory pathway is known to affect circadian period,^41^ the other negative clock regulatory pathway, casein kinase I (CKI)-PER, has been linked to period length as well.^24^ To evaluate its period effects, we used a CKIδ/ε-selective inhibitor, PF-670462, which also showed a dose-dependent period-lengthening effect on *Bmal1* in U2OS cells (**Figs. 2C, 2D, S3,** and **Table S1)**. It has been reported that PF-670462 has an IC50 of 0.55 μM for CKIε and an IC50 of 0.14 μM for CKIδ in whole cells.^42^ This molecule is proposed to increase circadian periods by specifically inhibiting CKIδ-mediated phosphorylation of PER, but not through CKIε.^34, 42^ We also ascertained the molecules’ abilities to lengthen the periods of *Per2*, via U2OS-*Per2:luc* reporter cells (**Fig. S4** and **Table S2)**. As anticipated, both compounds significantly increased the period of *Per2;* period enhancements were of similar magnitude in both reporter cell lines (**Fig. S5**).

**Figure 2.**
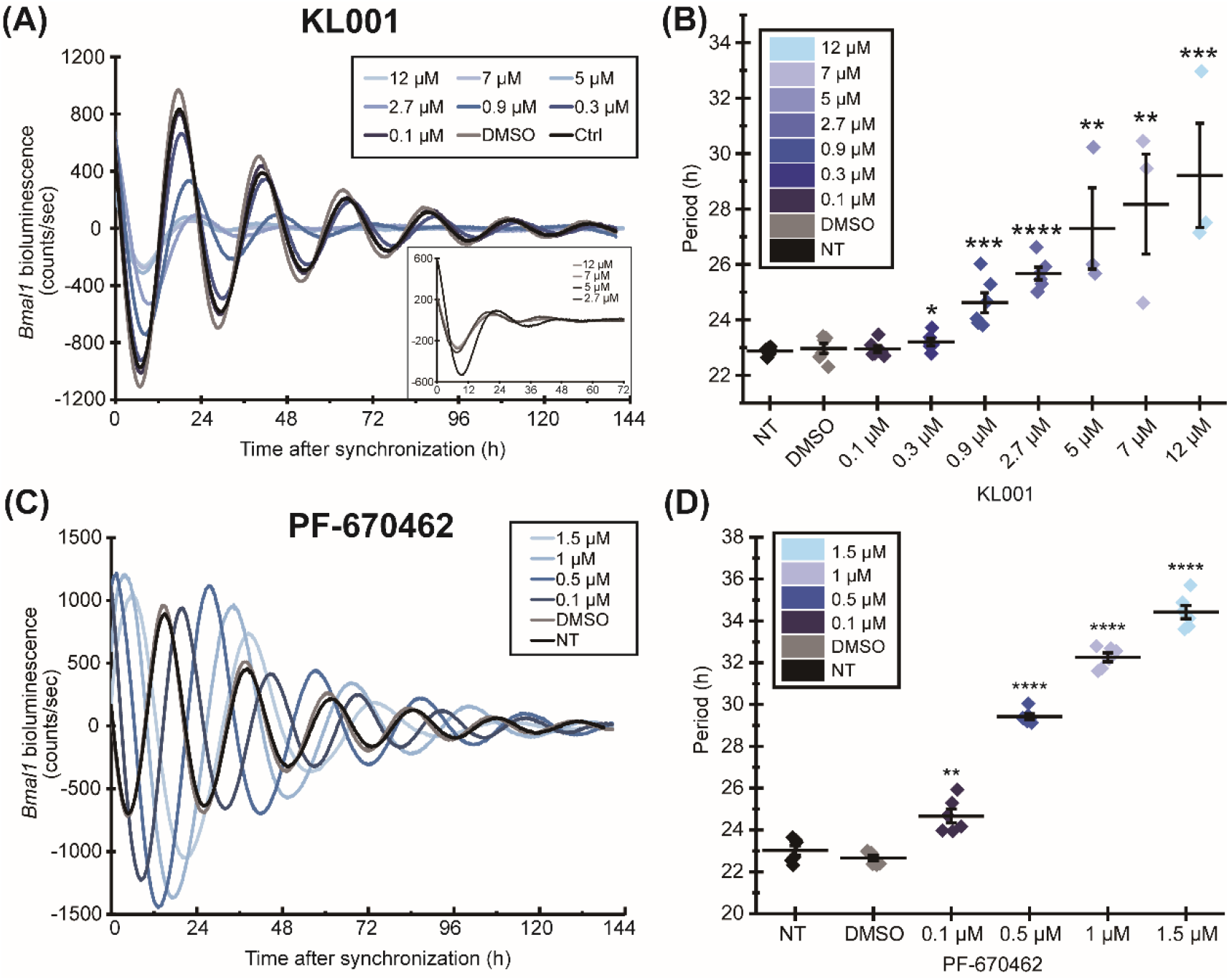
Direct circadian modulators KL001 (**A,B)** and PF-670462 (**C,D)** increase the circadian period of *Bmal1* in a dose-dependent manner. The inset in (**A)** shows an enlarged version of the first three cycles for treatments at the three highest concentrations. Representative traces are shown; replicate traces (raw and detrended) for each condition are in **Fig. S2** and **S3**. The line represents the mean of the data±1 SE. Statistical significance was evaluated via paired student T-test, * p<0.05, ** p<0.01, *** p<0.001, and ****p<0.0001. NT = non-treated; DMSO = DMSO-only control (0.2%). Period and goodness of fit (GOF) for each condition is shown in **Table S1**.

The indirect circadian-affecting molecule SP600125 (a JNK inhibitor with a cell-free IC50 of 40 nM for JNK1 and 2, and 90 nM for JNK3)^43^ also increased *Bmal1* periods in a dose-dependent manner (Fig. 3A**, 3B, S6,** and **Table S3**). However, while we observed that other molecules had similar period enhancements for *Bmal1* and *Per2*, SP600125 had a greater effect on *Bmal1* than *Per2* periods (**Fig. S5, S7,** and **S8)**. It has been shown that activated JNK isomers can phosphorylate BMAL1-CLOCK complexes to regulate the speed and phase of circadian oscillations in cells.^44^ Hence, SP600125 may lengthen the period via activation of JNK and through its downstream targets, including c-jun and AP-1, which regulate *Per1* and *Per2* transcriptional activities and suppress glucocorticoid receptor activity (critical for resetting circadian time).^35, 45^ However, evidence also exists that this molecule may alternatively target CKI, which phosphorylates PER.^40^

**Figure 3.**
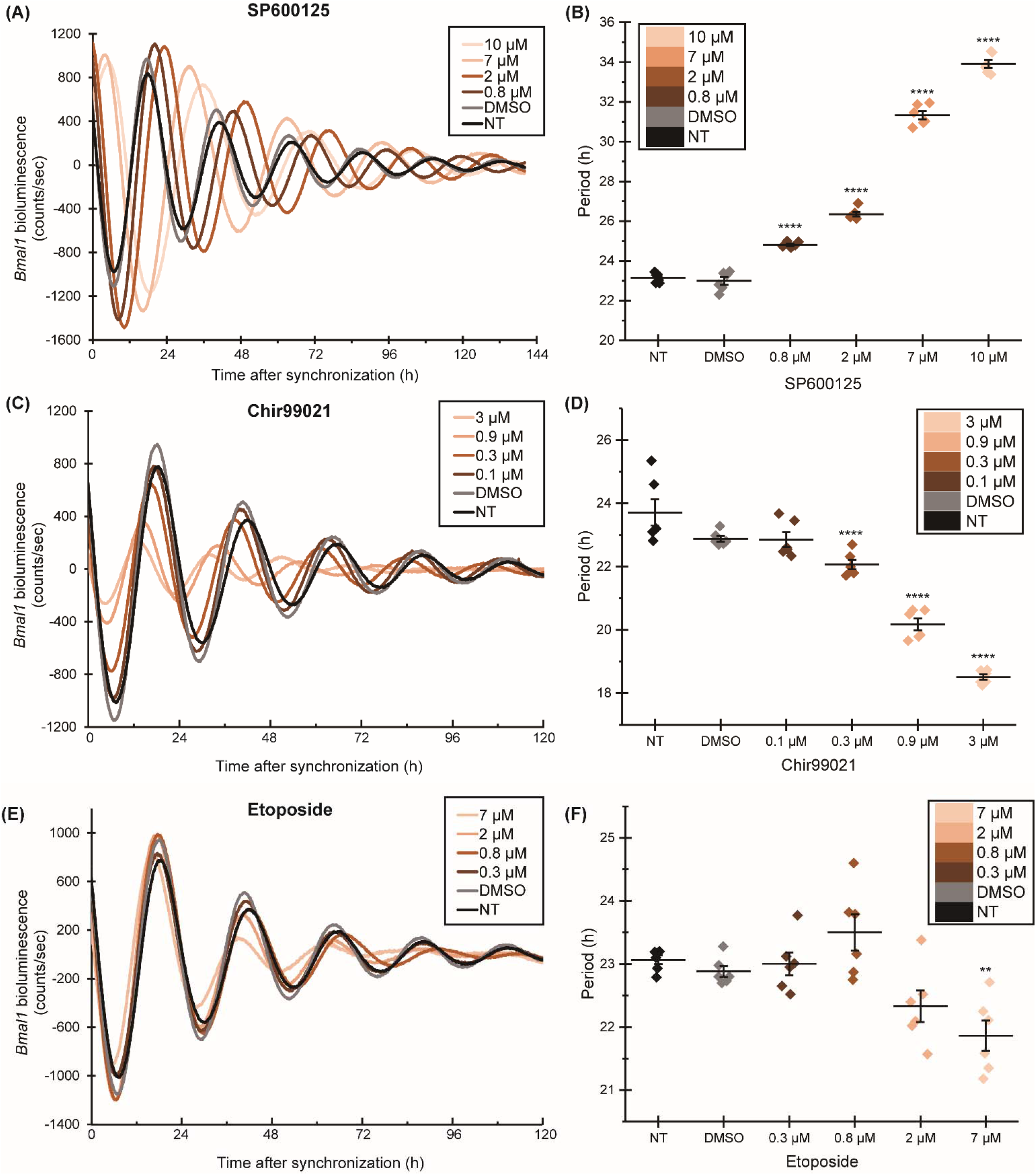
Three indirect circadian modulators, SP600125 (**A,B)**, Chir99021 (**C,D)** and Etoposide (**E,F)**, significantly increase or decrease the period of *Bmal1* in U2OS cells. Representative traces are shown; replicate traces (raw and detrended) for each condition are in **Figs. S7, S9,** and **S10**. The line represents the mean of the data±1 SE. Statistical significance was evaluated via paired student T-test, ** p<0.01 and **** p<0.0001. NT = non-treated; DMSO = DMSO-only control (0.2%). Individual period and goodness of fit (GOF) for each drug treatment are shown in **Table S3**.

While KL001, PF-670462, and SP600125 all displayed period-lengthening effects, Chir99021 (a GSK3α/β inhibitor with a cell-free IC50 of 10 nM for GSK3α and 6.7 nM for GSK3β)^46^ and Etoposide (a TOPII inhibitor with cell type-dependent IC50 ranges from 0.1 to >100 µM) shortened periods. Chir99021 significantly decreased *Bmal1* periods in a dose-dependent manner (Fig. 3C**, 3D, S9,** and **Table S3)**, however, Etoposide showed significant reduction of period only at the highest concentration tested (Fig. 3E**, 3F, S10,** and **Table S3)**. Both molecules showed similar effects on *Per2* transcription (**Fig. S7, S8)**. Chir99021 specifically inhibits GSK3α/β, which prevents phosphorylation of PER2,^47^ CRY2,^48^ CLOCK,^49^ BMAL1,^32^ and REV-ERVα.^50^ It has been shown that its use in different cell models results in short circadian periods. On the other hand, Etoposide may reduce period by interacting with topoisomerize type II (TOPII), which has been identified as a regulator of *Bmal1* transcriptional activity.^51^ Taken together, all five small molecules demonstrated an ability to modulate circadian periods of *Bmal1* and *Per2* in dose-dependent manners not attributable to toxicity (**Fig. S11).**

### Stabilization of PER and CRY have distinct effects on cancer cell migration

CRY proteins are generally characterized as tumor suppressors.^52^ Ye et al. recently reported a comprehensive analysis of 51 clock and clock-controlled genes across 32 cancer types, finding that 90.2% of them are either up- or down-regulated in at least one cancer.^53^ Specifically, CRY2 was found to be down-regulated in most cancers, and its suppression was associated with worse patient survival rates. The cancer suppressive role of CRY2 was also evaluated by Huber et al., whose mechanistic data indicated that CRY2 and FBXL3 cooperatively destabilized c-MYC (a critical oncoprotein for cancer cell proliferation), and disruption of *Cry2* may contribute to elevated c-MYC and higher cancer susceptibility in shift-workers and animals.^54^ However, Chun et al. found that a direct CRY inhibitor, KS15, significantly inhibited breast cancer cell proliferation and increased drug responses to chemotherapies.^55^

Given that CRY is down-regulated in various cancer types, it was hypothesized that stabilization of CRY via KL001 treatment might lead to reduction of cellular metastatic characteristics. A primary feature of metastasis is the migration of primary cancer cells to other, secondary sites. To assess whether CRY stabilization may influence cancer cell motility, U2OS cells were treated with KL001 and cell migration was evaluated via wound healing assays. Unexpectedly, our results showed that higher concentrations of KL001 yielded slightly increased cancer cell migration at early time points following treatment (T=6, 12, and 18 h), but resulted in no significant overall (24 h) change (Fig. 4A). However, it is likely that changes occur in a cell-dependent manner; Xue et al. used KL001 to impair mitosis and epithelium repair in corneal wound healing.^56^

**Figure 4.**
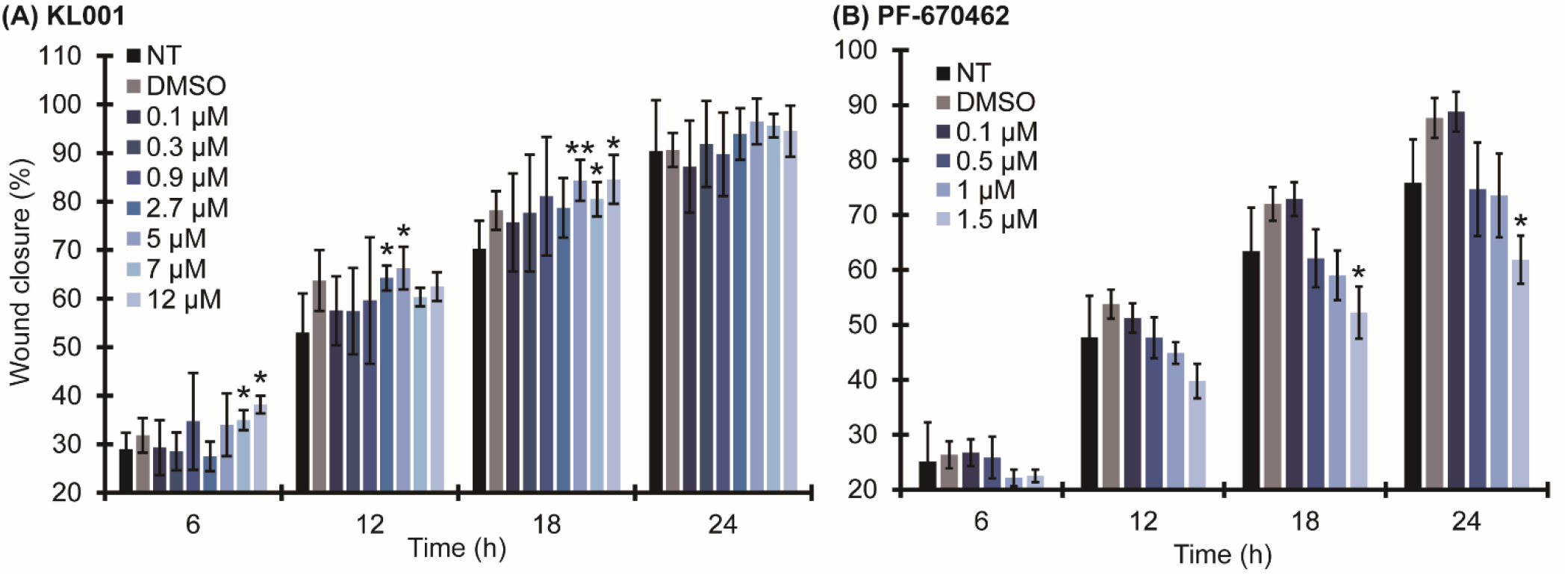
Direct circadian modulators KL001 (**A)** and PF-670462 (**B)** alter the motility of U2OS-*Bmal1:luc* cells. 5, 7, and 12 μM KL001 promote cell motility at T=6, 12, and 18 h (**A)**. 1.5 μM PF-670462 reduces cell motility (**B)**. Data are representative of three biological replicates; error bars represent standard deviation. Statistical significance was evaluated via paired student T-test, * p<0.05 and ** p<0.01. NT = non-treated; DMSO = DMSO-only control (0.2%).

Concurrently, we found that 1.5 μM PF-670462 treatment significantly inhibited cell motility (Fig. 4B). Its targets, CKIε and δ, are key regulators for many biological processes, including circadian rhythms, cell cycle, cell adhesion, cytoskeleton construction, and receptor-coupled signal transduction.^57^ Meng et al. showed that selective inhibition of CKIδ via PF-670462 stabilized the nuclear localization of endogenous PER2 in both SCN and peripheral cells, indicating that the molecule can act as a PER stabilizer.^34^ Our results in Fig. 4 suggest that stabilization of CRY and PER may influence cancer cell migration differently. However, other mechanistic roles of CKI need to be taken into consideration as well. CKI has been implicated in numerous oncogenic signaling networks, including Wnt/β-catenin, p53, PI3K/AKT, NFκB, TGFβ, and Hedgehog pathways, and contradictory evidence has implied that the CKI family may play dual-roles as tumor drivers and suppressors.^57, 58^ While we observed that PF-670462 inhibited osteosarcoma cell migration (Fig. 4B), the same treatments resulted in promotion of cell proliferation (**Fig. S12**). Although different mechanisms for regulating cell proliferation and migration via CKIε/δ activity may exist, we suspect that inhibition of migration via PF-670462 was induced through the TGFβ pathway: it has been shown that this molecule can prevent TGFβ-induced epithelial-mesenchymal transitions that result in enhanced motility.^59^

### Circadian period length may not be correlated with cell migration

Many cellular functions, including cell cycle, proliferation, and migration, are controlled by circadian rhythms, but little is known regarding the association between period length and the rates of these processes. As described above, the two direct circadian modulators, KL001 and PF-670462, resulted in opposite outcomes in their effects on cancer cell migration, although they both increased *Bmal1* and *Per2* periods in U2OS cells. Differing results were also observed between the two indirect circadian modulators with period-shortening effects: Chir99021 suppressed cell migration in a dose-dependent manner, while Etoposide showed no significant change (Fig. 5). As described above, Chir99021 targets GSK3α and β (serine/threonine kinases), which play important roles in many chronic progressive diseases, including diabetes mellitus, neurodegenerative disorders, and cancer.^60^ While GSK3β is a key regulator in the Wnt/β-catenin pathway,^61^ no data exists to rationalize how activation of β-catenin may result in lower cell motility. However, it has been proposed by Kobayashi et al. that the activities of GSK3α and β are tightly associated with cell migration, and another GSK3 inhibitor suppressed cellular migration through down-regulation of the focal adhesion kinase (FAK)/guanine nucleotide exchange factors (GEF)/Rac1 pathway.^62^ Our results may be indicative of this route (Fig. 5A).

**Figure 5.**
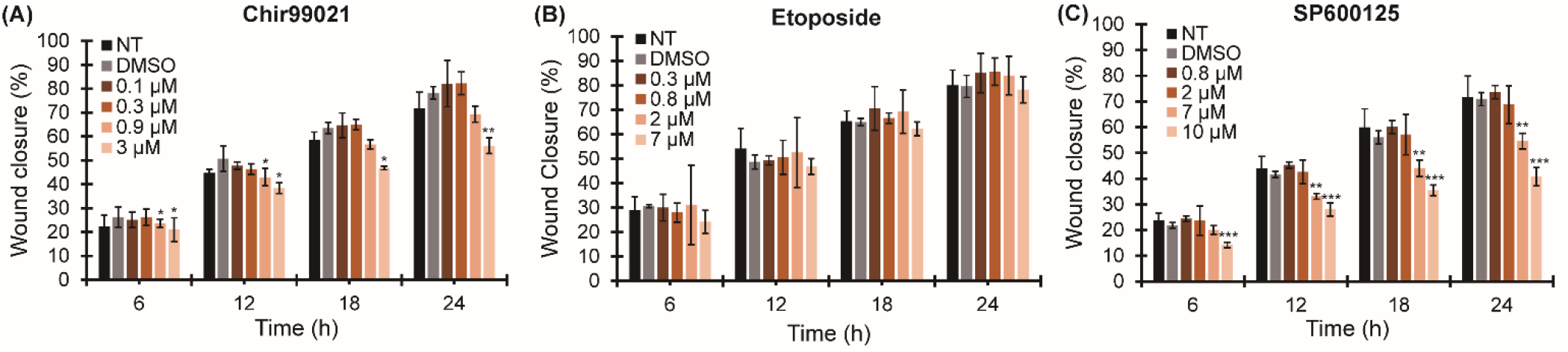
Three indirect circadian modulators, Chir99021 (**A)**, Etoposide (**B)**, and SP600125 (**C)**, influence cell motility to various levels. SP600125 and Chir99021 treatments significantly inhibit cell migration (**A,C)**, while Etoposide shows no effect (**B)** in U2OS cells. Data are (**D)** representative of three biological replicates; error bars represent standard deviation. Statistical significance was evaluated via paired student T-test, * p<0.05, ** p<0.01, and *** p<0.001. NT = non-treated; DMSO-only control (0.2%).

Unlike Chir99021, Etoposide did not affect cell migration (Fig. 5B) or viability (**Fig. S11E)**. The target of Etoposide is TOPII, which is known to regulate cell proliferation. There are two TOPII isoenzymes that exist in mammals, TOPII-α and TOPII-β.^63^ TOPII-α is a proliferation marker overexpressed in tumor cells, whereas TOPII-β is expressed in proliferating and post-mitotic cells.^64^ While Etoposide targets both TOPII isoenzymes, the relative contributions of TOPII-α and TOPII-β to its chemotherapeutic effects have yet to be resolved. Additionally, no data exists to demonstrate a connection between TOPII and cell migration. Thus, it is not surprising that Etoposide-mediated inhibition of TOPII in U2OS cells does not affect their migratory behavior. This result also implies that TOPII may not be involved in any pathways that could affect cell motility.

Another indirect circadian modulator, SP600125, increased circadian periods in U2OS cells and also inhibited cell migration (Fig. 5C). One possible mechanism to explain the inhibition of cellular motility resulting from SP600125 treatment is blockage of the JNK-matrix metalloproteinase (MMP) signaling cascade.^65^ The activity of MMPs, especially MMP-2 and −9, has been correlated with cancer cell invasion and migration. Our results are supported by the work of Fromigué et al., who inhibited upstream targets of JNK via atorvastatin treatment, and showed downregulation of MMPs, which resulted in reduction of osteosarcoma cell invasion and migration.^65^

Taken together, these data appear to indicate that circadian period length may not be correlated with cell migration, and that the pharmacological targets may play a dominant role in controlling cell motility. However, it is notable that PF-670462, SP600125, and Chir99021 affected circadian periods to greater extents (1.5 μM PF-670462 increased *Bmal1* period to 34.4± 0.8 h, 10 μM SP600125 increased *Bmal1* period to 33.9±0.5 h, and 3 μM Chir99021 decreased *Bmal1* period to 18.5±0.2 h), while KL001 and Etoposide only resulted in minor changes (12 μM KL001 increased the period of *Bmal1* to 29.2±3.3 h and 7 μM Etoposide decreased it to 21.9±0.6 h). The ability to alter circadian period to a greater extent may contribute to the repressive effects on cell migration. To be certain, identification and assessment of additional small molecules directly targeting core clock proteins are required.

### Effects of small molecules on formation of cancerous colonies

While several of the molecules studied here affected cancer cell migration, we wanted to assess whether they could influence other oncogenic features, such as tumor formation and proliferation. We evaluated effects on cell proliferation in a 3D environment, via soft agar assay. By assessing cancer cell proliferation in these semi-solid matrices, we are able to mimic what may occur in *in vivo* cellular microenvironments. After four weeks of incubation, U2OS cancer cells generally form “tumor-like” colonies in the agar gels. When cultured with the highest concentrations of each drug used elsewhere in this study, we observed that 1.5 μM PF-670462 significantly increased colony numbers (Fig. 6A), consistent with 2D proliferation (**Fig. S12B)**. As mentioned above, the CKI family can play dual roles in many cancer types.^57, 58^ Our data provide support for this in that PF-670462 suppresses motility but promotes cell proliferation of osteosarcoma cells.

**Figure 6.**
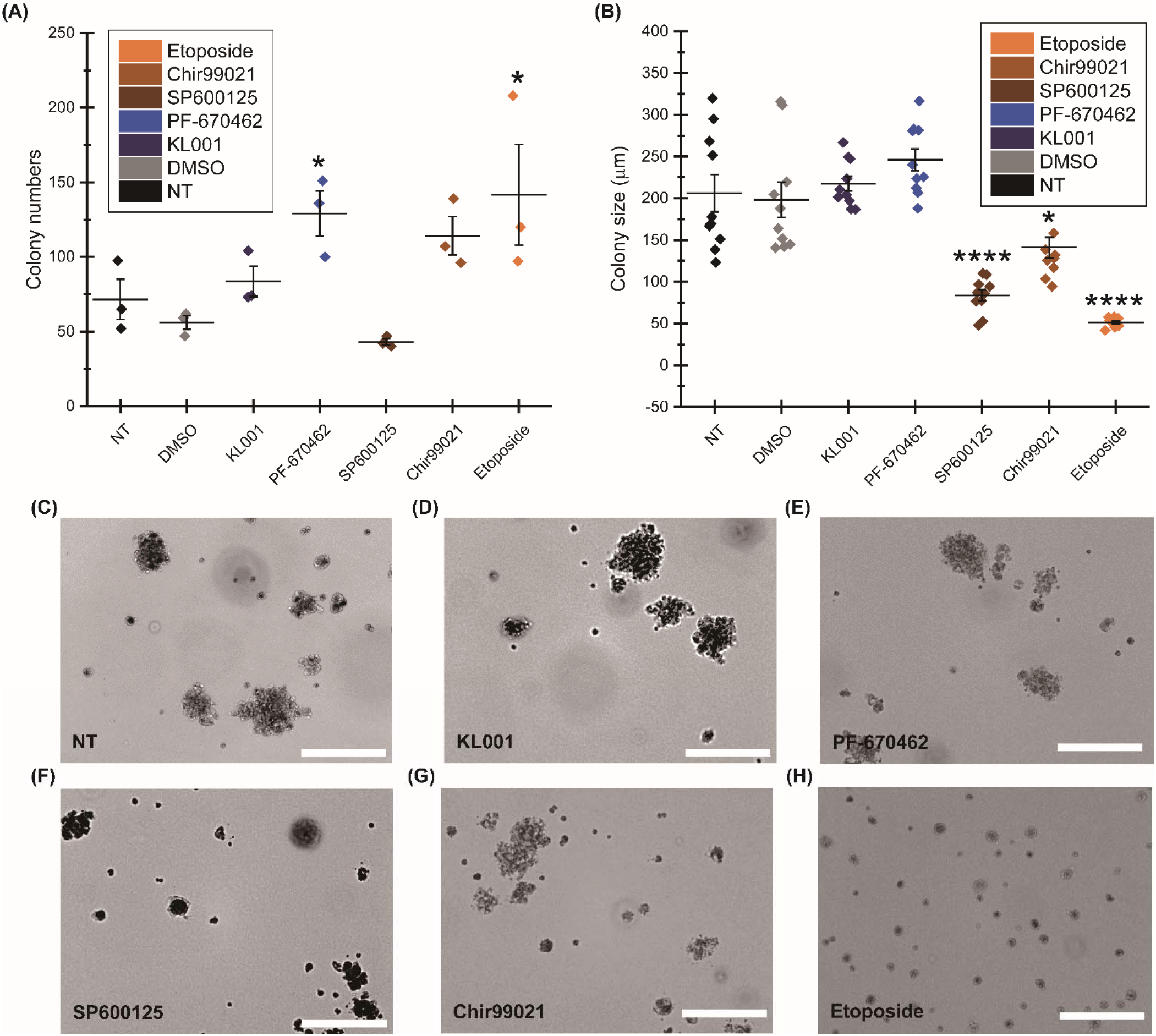
Effects of small molecules on tumor formation in a soft agar assay. After four weeks of incubation, colony numbers (**A)** and sizes (**B)** were evaluated. PF-670462 and Etoposide significantly increased colony numbers, while SP600125, Chir99021, and Etoposide reduced colony sizes. Statistical significance was evaluated via paired student T-test, * p<0.05 and **** p<0.0001. NT = non-treated; DMSO = DMSO-only control (0.2%). Representative images for various treatments are shown (**C-H)**. Scale bars are 100 μm.

Concurrently, treatment of 7 μM Etoposide significantly decreased colony size by 75%, from 206±70 μm to 51±5 μm (Fig. 6B). We suspect this was related to inhibition of cancer cell proliferation, not observed with treatment of Chir99021 (**Figs. S13A** and **S13B)**. Etoposide likely resulted in higher colony numbers on account of its cancer cell and colony growth suppression, which resulted in a lack of aggregation forming clusters (Fig. 6A). Inhibition of cell proliferation via Etoposide treatment has been observed previously.^66^ Additionally, treatment with 10 μM SP600125 and 3 μM Chir99021 decreased colony sizes by 59% and 32%, respectively, but had no effects on numbers of colonies relative to non-treated controls. This data indicates that Chir99021 and SP600125 may inhibit cell proliferation in a 3D environment (Fig. 6), but not in 2D monolayer cultures (**Figs. S13B** and **S13C)**. This is not surprising given that it is well-established that cells in two-versus three-dimensions may respond differently to drugs,^67^ underscoring the importance of this type of assessment.

## Conclusion

Disruption of circadian rhythms has been shown to elicit pathological development associated with cancer and other diseases. While many small molecules have been developed to modulate circadian rhythms, a systematic evaluation that uses the same concentration ranges in a parallel comparison of their effects on cellular and oncogenic characteristics is lacking. Here, five small molecules were used to modulate circadian period (e.g. increase or decrease the period of the circadian cycle) in the human osteosarcoma U2OS cell line. We found that they can influence periods and cancer cell migration, proliferation, and tumor formation to varying extents. KL001 and PF-670462 are two direct circadian modulators, which were previously shown to stabilize CRY and PER respectively. While both significantly increased the periods of *Bmal1* and *Per2* in U2OS cells, they exhibited opposing effects on cell migration. At T=6, 12, and 18 h, KL001 promoted cancer cell migration, but had no effects on cell proliferation or colony formation. Conversely, PF-670462 inhibited cell migration at the highest concentration tested, and enhanced cell proliferation and colony numbers.

SP600125, Chir99021, and Etoposide are three indirect circadian modulators, all of which significantly altered the periods of *Bmal1* and *Per2* in U2OS cells. Treatment with SP600125 increased periods, while Chir99021 and Etoposide resulted in decreases. In evaluating their effects on cancer cell behaviors, we hypothesized that we would observe opposing trends between period-lengthening and period-shortening groups in altering cell migration and colony formation. However, we found that both SP600125 and Chir99021 treatments inhibited cell migration, and resulted in diminished colony sizes. On the other hand, Etoposide showed no effect on cell migration but significantly reduced cell proliferation and colony size, which may be attributed to inhibition of TOPII, a key enzyme in DNA cleavage/ligation and cell proliferation regulation, likely not involved in cell migratory pathways. These data indicate that circadian period length may not be correlated with cell growth and migration.

Our data suggest that directly targeting circadian proteins or inducing circadian period effects may not be sufficient to modulate cancerous phenotypes. It is plausible that non-circadian pathways have a greater influence on cellular behavior. However, based on our results, it may also be possible that cellular effects are only observed upon significant deviation from the twenty-four hour cycle. There is no shortage of work to be done in this area, including assessment of other circadian alterations, including amplitude and phase, use of additional small molecules directly targeting circadian proteins that have greater effects on period, utilization of next generation sequencing (NGS) methods to further assess connections between small molecule-affected cancer and circadian pathways, and employment of combined genetic and chemical biology approaches via knockdown of non-circadian targets and chemical modulator treatments to further distinguish circadian-based from alternative pathway-based effects. Understanding the contributions of circadian rhythms and their effects on cellular phenotypes are critical to understanding roles of the clock in homeostasis and disease.

## Author Contributions

The manuscript was written by Lin, H.-H. and revised by Farkas, M. E. All authors have given approval to the final version of the manuscript.

## ACKNOWLEDGMENT

H.-H. L. was supported by a University of Massachusetts Amherst Chemistry-Biology Interface (CBI) training fellowship. K. L. R. was supported by a Summer Undergraduate Research Fellowship from the American Chemical Society. We gratefully acknowledge the laboratories of Prof. Yubing Sun (Mechanical Engineering, UMass Amherst) and Prof. Jungwoo Lee (Chemical Engineering, UMass Amherst) for plate reader use, and Prof. D. Joseph Jerry (Veterinary and Animal Sciences, UMass Amherst) for advice on stable transfections.

