## Supporting Information for "Oncogenic and Circadian Effects of Small Molecules Directly and Indirectly Targeting the Core Circadian Clock"

**Methods**

**Luciferase Assay** Cells were seeded in 24-well plates with 500 μL of 2 x 10^5^ cells/mL. After cells reached 100% confluence, they were lysed and assessed using the Luciferase Assay System (Promega) according to the manufacturer’s instructions. Luminescence from U2OS-*Bmal1:luc* cells was measured via a SpectraMax M5 multi-mode microplate reader and luminescence from U2OS-*Per2:luc* cells was measured via a Biotek Syngery H1 multi-mode microplate reader.

**Cell Viability Assay** Cells were seeded in 24-well plates (Nunclon) with 500 μL of 2 x 10^5^ cells/mL and incubated for approximately 24 h until they reached 100% confluence. The cells were then treated with indicated small molecules and incubated for 24 h, at which time the culture media was replaced with 400 μL/well of 10% Alamar blue dye (Invitrogen). The cells were incubated with the dye at 37 °C with 5% CO_2_ for 1 h, and 200 μL of solution from each well was transferred to a black 96 well plate (Costar). Samples were exposed to an excitation wavelength of 560 nm and tracking emission wavelength of 590 nm, using a Biotek Syngery H1 multi-mode microplate reader.

**Cell Proliferation Assay** Cells were seeded in 24-well plates with 500 μL of 1 x 10^5^ cells/mL. After 24 h incubation, the cells were approximately 50% confluent. They were then dosed with 400 μL/well of the designated small molecules at the indicated concentrations in 10% Alamar blue. The first time point (T=0) was taken immediately after drug treatment, with subsequent analyses every 24 h thereafter until 96 h. At every time point, 200 μL of solution was transferred from each well to a black 96 well plate, and exposed to an excitation wavelength of 560 nm, tracking emission wavelength of 590 nm, using a Biotek Syngery H1 multi-mode microplate reader.

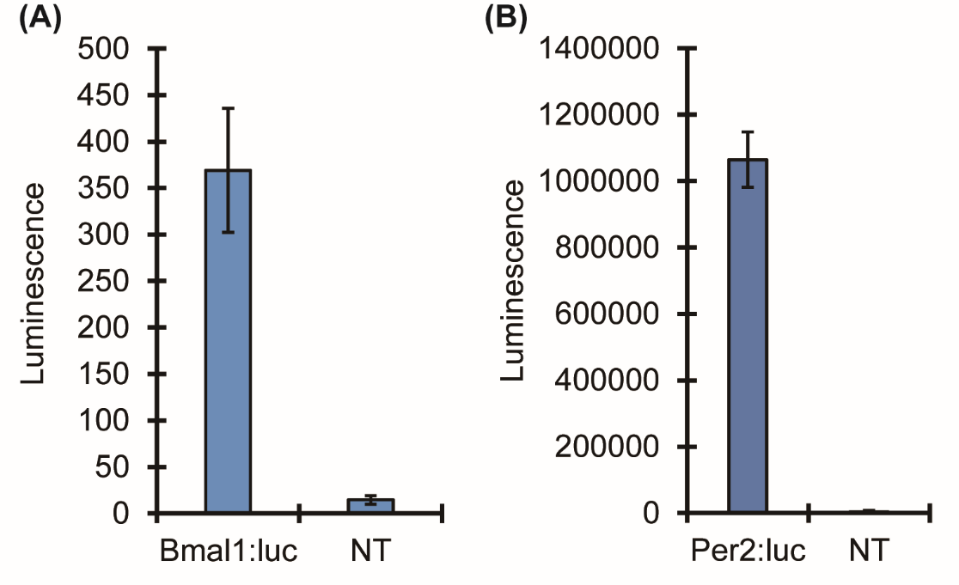

**Figure S1.** U2OS cells were stably transfected with *Bmal1:luc* and *Per2:luc* reporters to facilitate the real-time tracking of circadian rhythms. The luciferase assay showed that cells contained luciferase proteins after inserting *Bmal1:luc* **(A)** and *Per2:luc* **(B)** into the host genome via viral transduction. Data are representative of three technical replicates; error bars represent standard deviation. NT = non-treated.

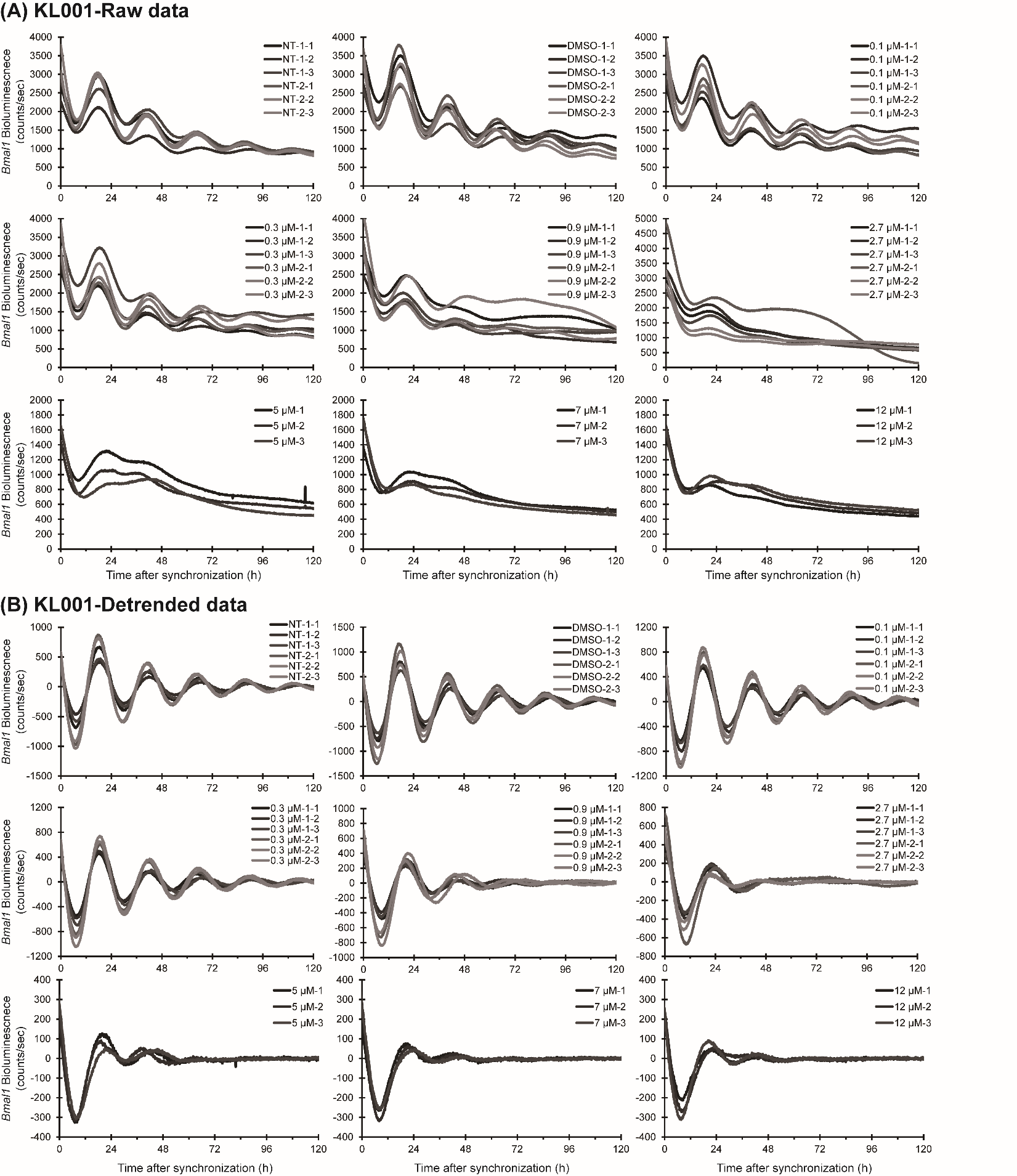

**Figure S2.** Individual luminometry oscillation curves (n=3 for two separate experiments, designated 1 and 2, except for 5, 7, and 12 μM treatments) for KL001-treated U2OS-*Bmal1:luc* cells. Raw data **(A)** and detrended data **(B)** are shown**.** NT = non-treated; DMSO = DMSO-only control (0.2%).

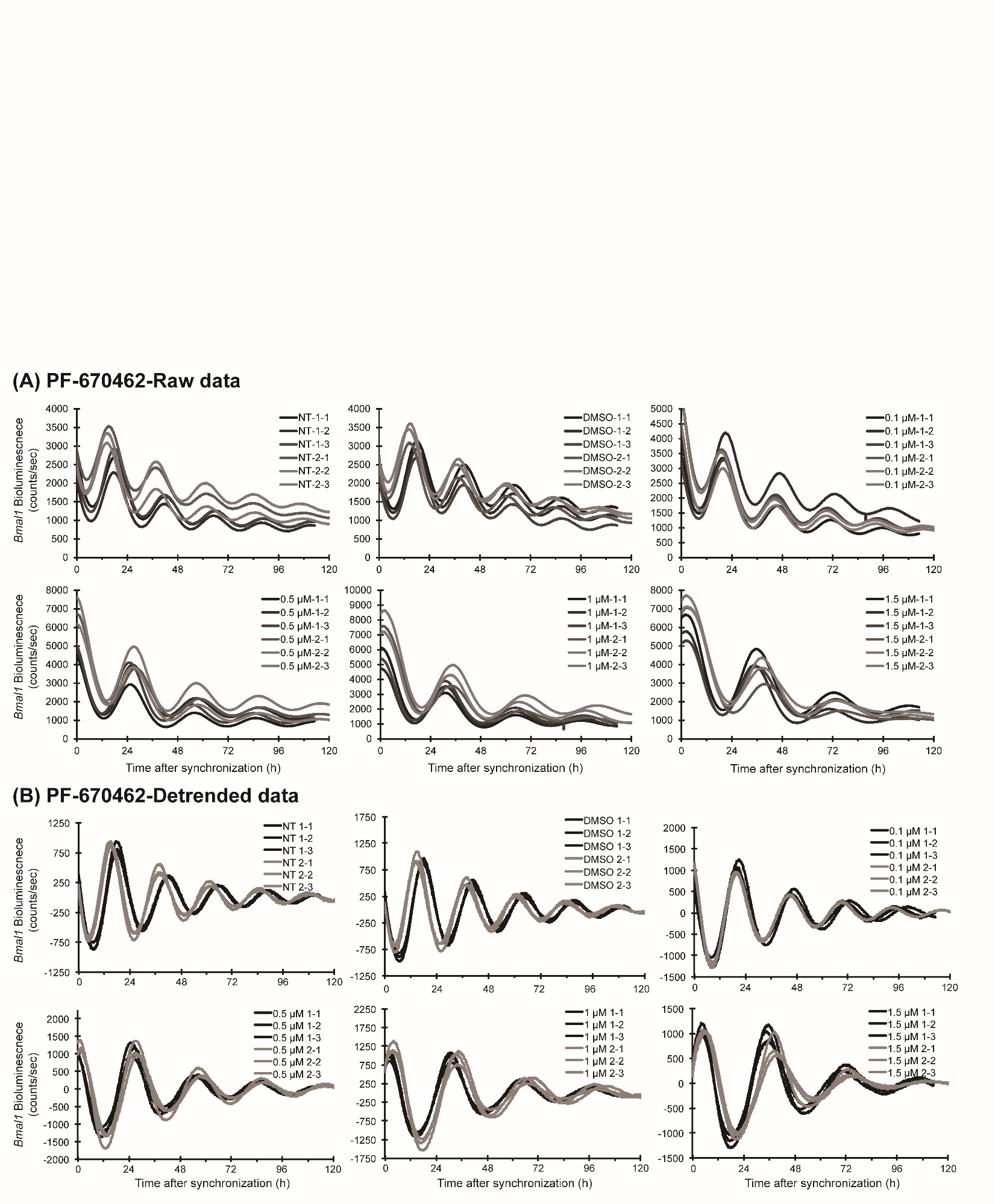

**Figure S3.** Individual luminometry oscillation curves (n=3 for two separate experiments, designated 1 and 2) for PF-670462-treated U2OS-*Bmal1:luc* cells. Raw data **(A)** and detrended data **(B)** are shown. NT = non-treated; DMSO = DMSO-only control (0.2%).

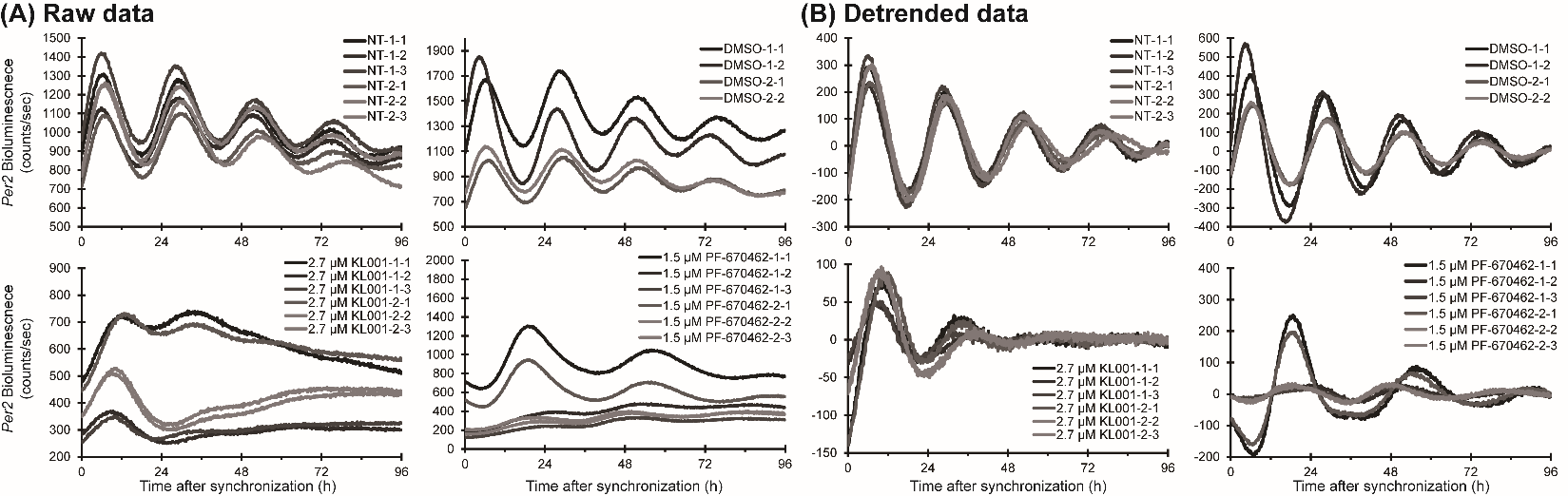

**Figure S4.** Individual luminometry oscillation curves (n=3 for two separate experiments, designated 1 and 2) for 2.7 μM KL001 and 1.5 μM PF-670462 treatments and controls in U2OS-*Per2:luc* cells. Raw data **(A)** and detrended data **(B)** are shown. NT = non-treated; DMSO = DMSO-only control (0.2%).

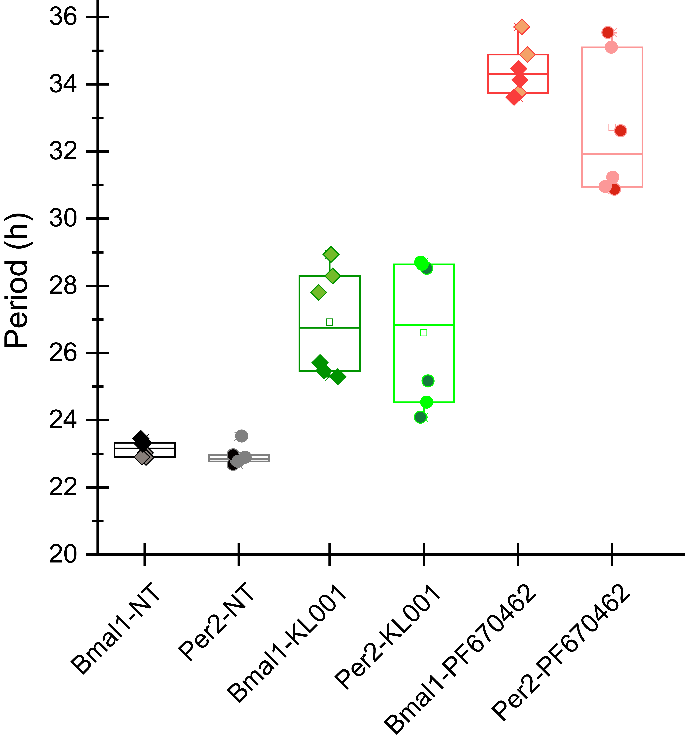

**Figure S5.** Comparison of maximum changes in period for U2OS-*Bmal1:luc* and U2OS-*Per2:luc* cells following treatment with direct circadian modulators KL001 and PF-670462. Different shades of the same color represent data points from independent experiments. Boxes represent the interquartile range (25th to 75th percentile), and the lines bisecting them represent the median. The small square in the center is the mean, and whiskers indicate the 5th and 95th percentiles. NT = non-treated.

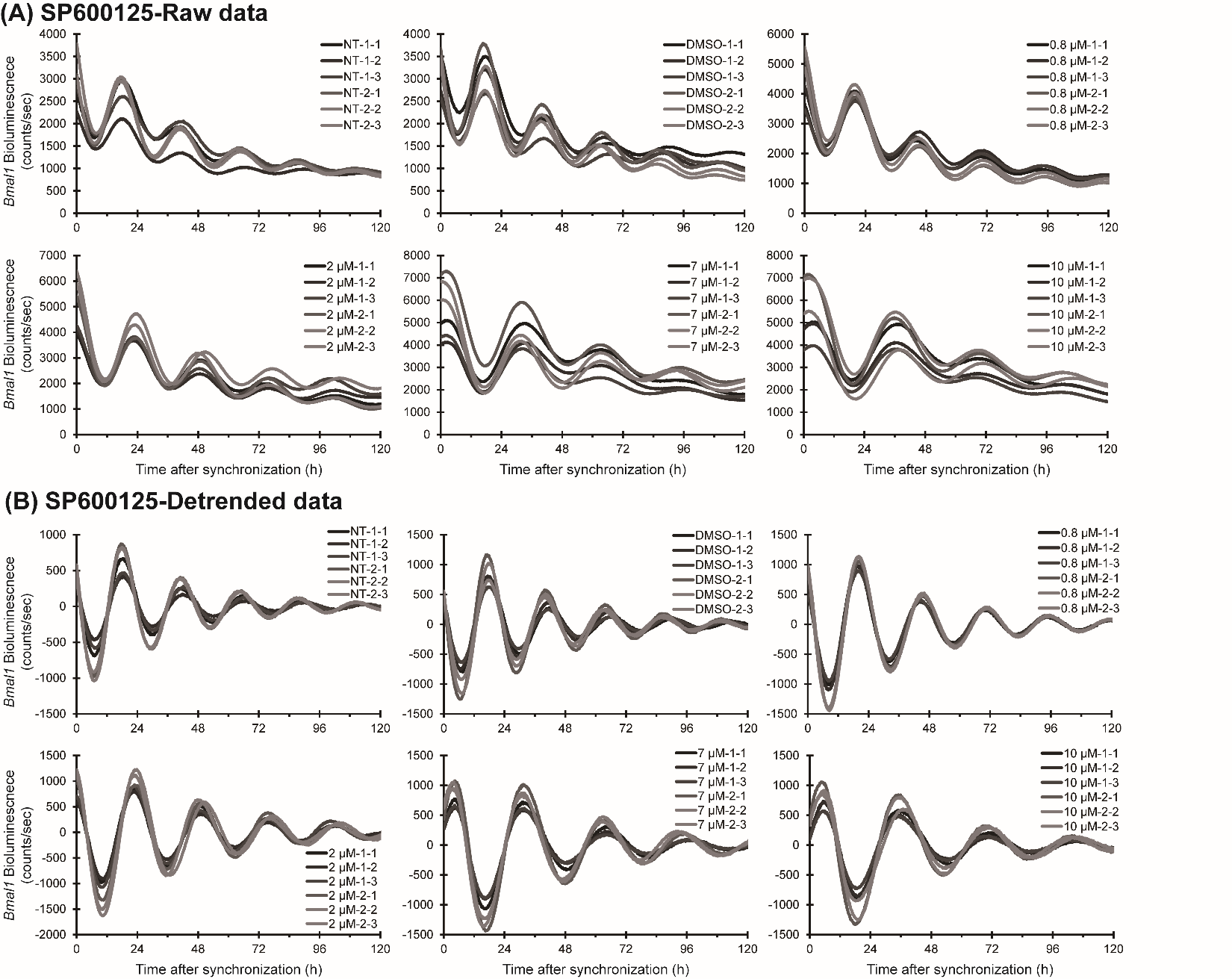

**Figure S6.** Individual luminometry oscillation curves (n=3 for two separate experiments, designated 1 and 2) for SP600125-treated U2OS-*Bmal1:luc* cells. Raw data **(A)** and detrended data **(B)** are shown. NT = non-treated; DMSO = DMSO-only control (0.2%).

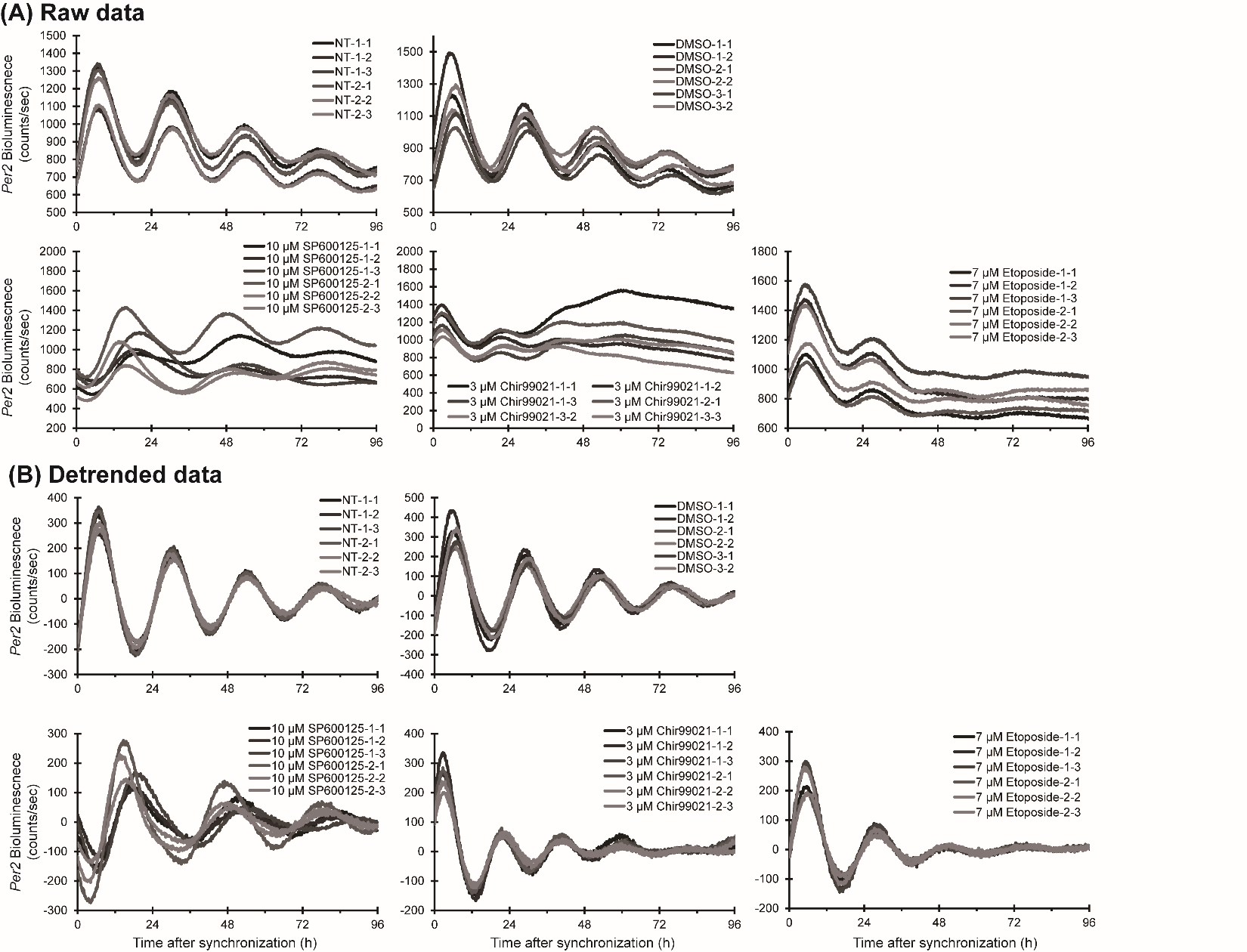

**Figure S7.** Individual luminometry oscillation curves (n=3 for two separate experiments, designated 1 and 2) for 10 μM SP600125, 3 μM Chir99021, and 7 μM Etoposide treatments and controls in U2OS-*Per2:luc* cells. Raw data **(A)** and detrended data **(B)** are shown. NT = non-treated; DMSO = DMSO-only control (0.2%).

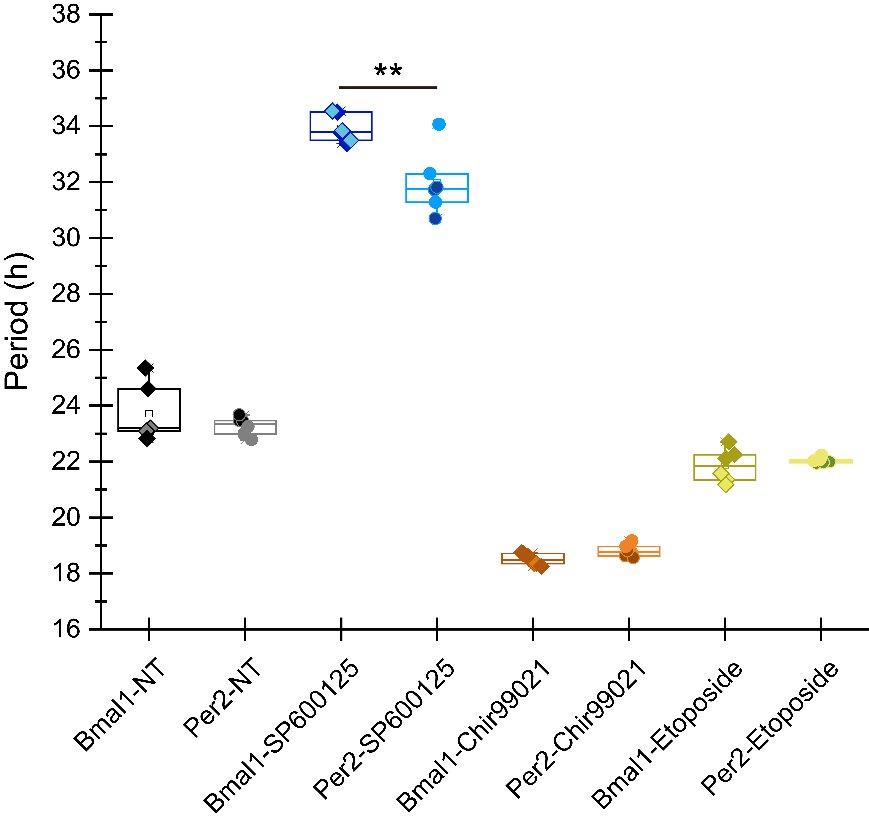

**Figure S8.** Comparison of changes in period for U2OS-*Bmal1:luc* versus U2OS-*Per2:luc* cells following treatment with indirect circadian modulators. Statistically significant differences were only observed in 10 μM SP600125-treated *Bmal1* and *Per2* cells, as determined by paired student T-test (** p<0.01). Different shades of the same color represent data points from independent experiments. Boxes represent the interquartile range (25th to 75th percentile), and the lines bisecting the boxes represent the median. The small square in the center of each box is the mean, and whiskers indicate the 5th and 95th percentiles. NT = non-treated.

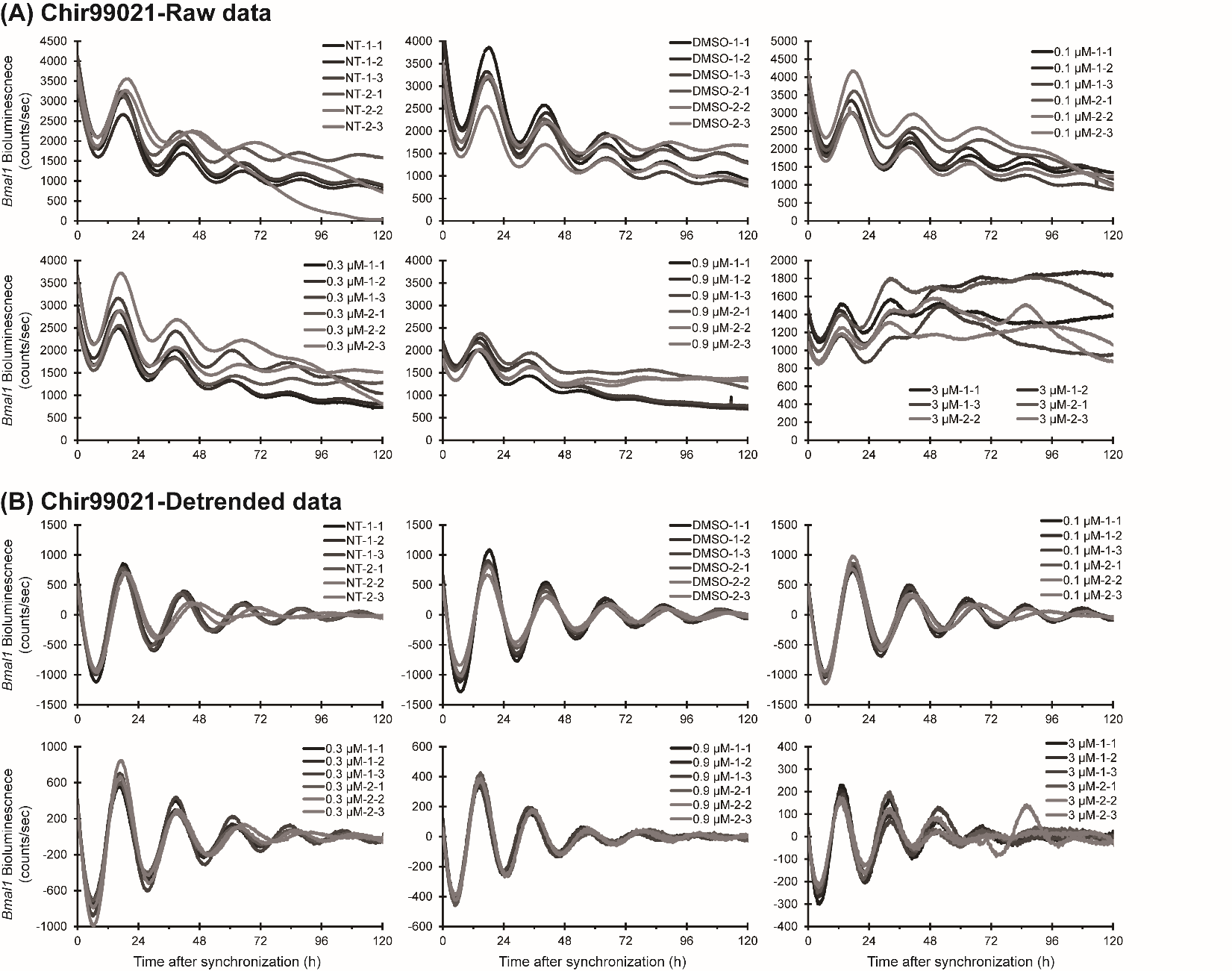

**Figure S9.** Individual luminometry oscillation curves (n=3 for two separate experiments, designated 1 and 2) for Chir99021-treated U2OS-*Bmal1:luc* cells. Raw data **(A)** and detrended data **(B)** are shown. NT = non-treated; DMSO = DMSO-only control (0.2%).

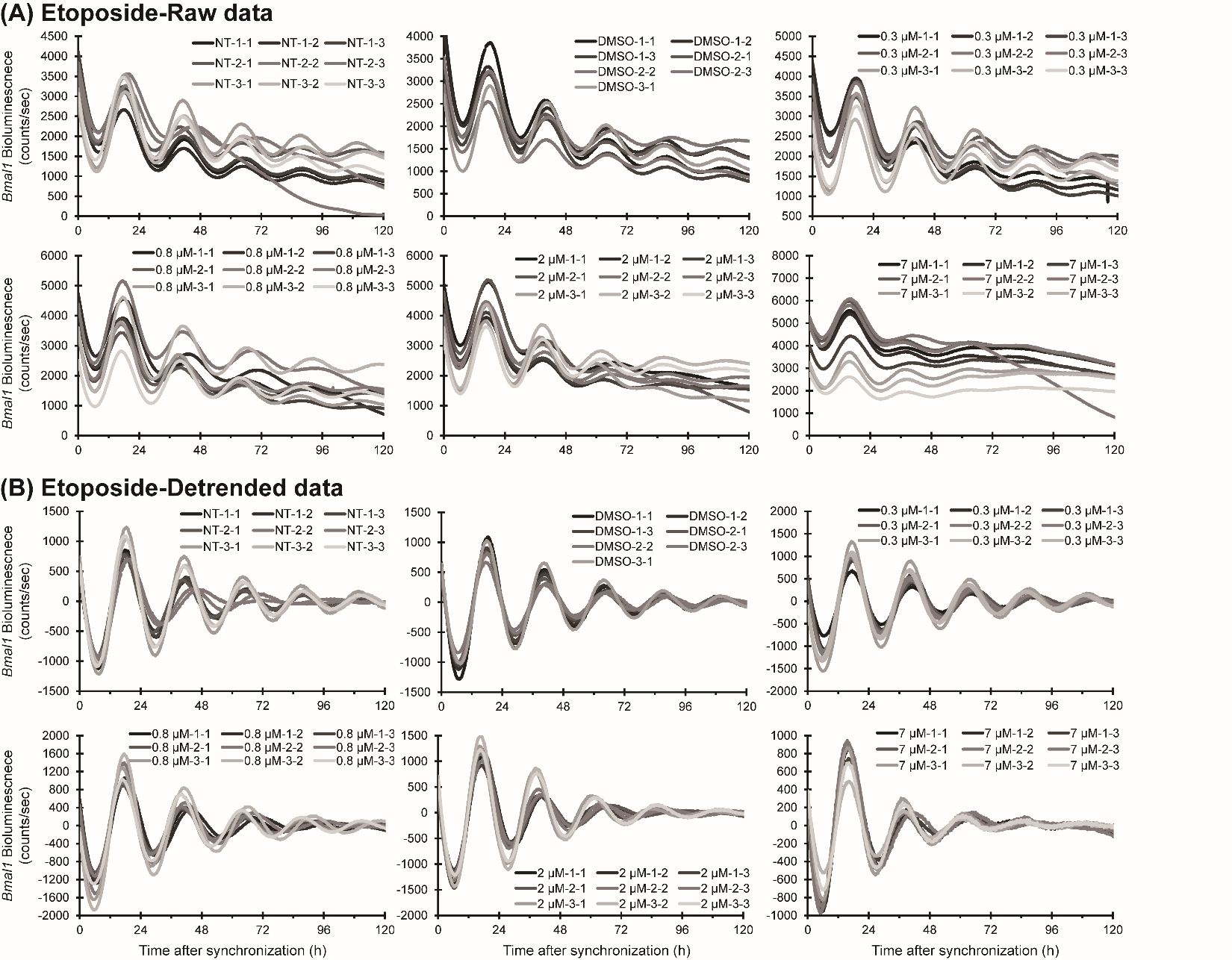

**Figure S10.** Individual luminometry oscillation curves (n=3 for three separate experiments, designated 1, 2, and 3) for Etoposide-treated U2OS-*Bmal1:luc* cells. Raw data **(A)** and detrended data **(B)** are shown. NT = non-treated; DMSO = DMSO-only control (0.2%).

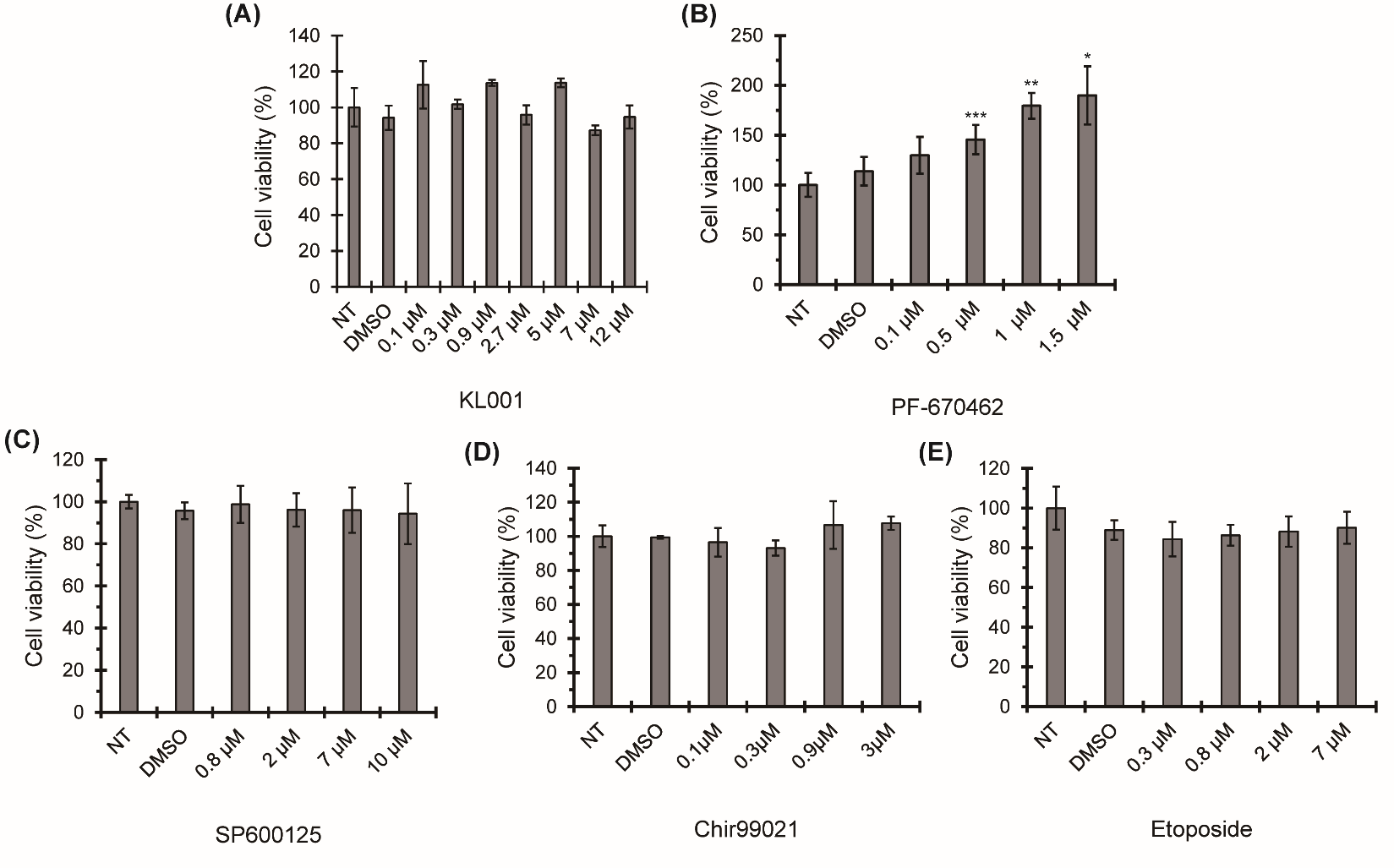

**Figure S11.** Viability assay with KL001 **(A)**, PF-670462 **(B)**, SP600125 **(C)**, Chir99021 **(D)**, and Etoposide **(E)** in U2OS cells. Cells were treated at indicated concentrations of the respective compounds for 24 h prior to AlamarBlue assay. No significant differences were observed between non-treated and KL001, SP600125, Chir99021, and Etoposide-treated cells as, determined by paired student T-test. PF-670462 showed increased cell viability, as corroborated by cell proliferation (**Fig. S12**) and colony formation (**Fig. 6**) assays. Error bars represent standard deviation across three biological replicates. Statistical significance was evaluated via paired student T-test, ***p<0.001, **p<0.01, *p<0.05. NT = non-treated; DMSO = DMSO-only control (0.2%).

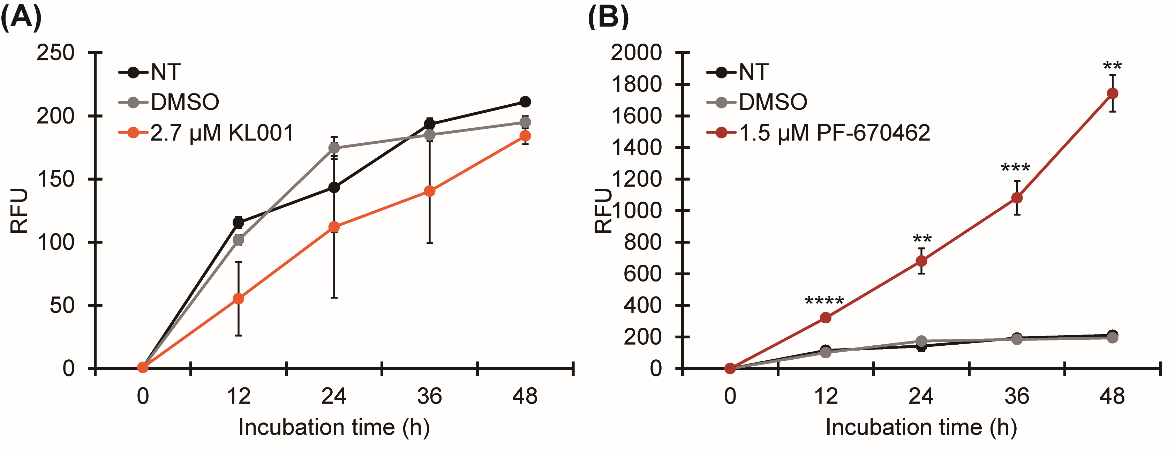

**Figure S12.** Proliferation assay with direct circadian modulators KL001 **(A)** and PF-670462 **(B)** in U2OS cells. 2.7 μM KL001 treatment did not affect cell proliferation in a statistically significant manner. 1.5 μM PF-670462 dramatically increased cell proliferation of U2OS cells, compared to non-treated or DMSO-treated cells. Error bars represent standard deviation across n=3 biological replicates. Statistical significance was evaluated via paired student T-test, **p<0.01, ***p<0.001, and ****p<0.0001. NT = non-treated, DMSO = DMSO-only control (0.2%).

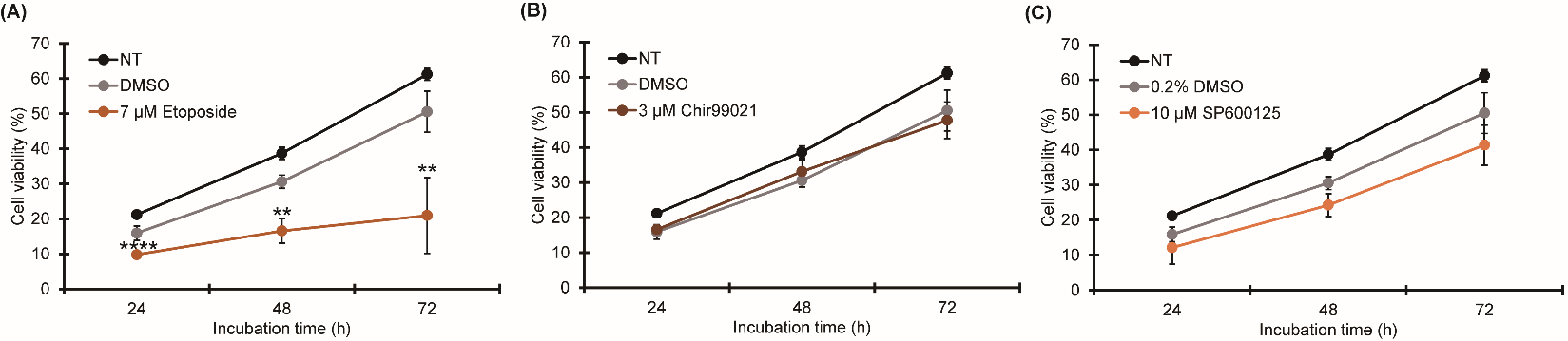

**Figure S13.** Proliferation assay with period-lengthening compounds, Etoposide **(A)**, Chir99021 **(B)**, and SP600125 **(C)** in U2OS cells. 10 μM SP600125 and 3 μM Chir99021 treatment did not affect proliferation, while 7 μM Etoposide significantly decreased cell proliferation of U2OS cells, compared to non-treated or DMSO-treated cells. Error bars represent standard deviation across n=3 biological replicates. Statistical significance was evaluated via paired student T-test, **p<0.01 and ****p<0.0001. NT = non-treated; DMSO = DMSO only control (0.2%).

**Table S1. Calculated periods and goodness of fits (GOFs) for oscillations of *Bmal1:luc* reporter signals in U2OS cells treated with direct circadian modulators and respective controls.**

| **KL001** | | | **NT** | **GOF (%)** | **DMSO** | **GOF (%)** | **0.1 μM** | **GOF (%)** | **0.3 μM** | **GOF (%)** | **0.9 μM** | **GOF (%)** | **2.7 μM** | **GOF (%)** |
| --- | --- | --- | --- | --- | --- | --- | --- | --- | --- | --- | --- | --- | --- | --- |
| **Exp 1** | | **replicate 1** | 23.03 | 99 | 22.81 | 98.3 | 22.86 | 96 | 23.05 | 99.1 | 24.66 | 98.8 | 28.93 | 95.1 |
|  |  | **replicate 2** | 22.89 | 98.8 | 22.66 | 98.6 | 22.79 | 98.5 | 23.11 | 99.2 | 24.04 | 98.3 | 28.29 | 96.3 |
|  |  | **replicate 3** | 22.9 | 98.9 | 22.31 | 98.6 | 23.09 | 98.7 | 23.37 | 98.9 | 26.02 | 98.5 | 27.8 | 95.4 |
| **Exp 2** | | **replicate 1** | 23.29 | 97.6 | 23.48 | 96.5 | 22.7 | 97.7 | 23.17 | 97.4 | 25.29 | 96.2 | 25.47 | 98.1 |
|  |  | **replicate 2** | 23.45 | 97.7 | 23.39 | 96.9 | 23.47 | 96.1 | 22.78 | 97.7 | 23.89 | 96.4 | 25.3 | 97.3 |
|  |  | **replicate 3** | 23.33 | 98.6 | 23.31 | 96.6 | 22.77 | 97.7 | 23.72 | 96.5 | 23.81 | 98.2 | 25.71 | 97.4 |
| **KL001** | | | **5 μM** | | **GOF (%)** | | **7 μM** | | **GOF (%)** | | **12 μM** | | **GOF (%)** | |
| **Exp 3** | **replicate 1** | | 26 | | 96.4 | | 24.61 | | 98 | | 27.51 | | 97.7 | |
|  | **replicate 2** | | 25.67 | | 79.1 | | 29.46 | | 97 | | 32.97 | | 89.1 | |
|  | **replicate 3** | | 30.23 | | 94.3 | | 30.45 | | 89.5 | | 27.15 | | 98.3 | |
| **PF-670462** | | | **NT** | **GOF (%)** | **DMSO** | **GOF (%)** | **0.1 μM** | **GOF (%)** | **0.5 μM** | **GOF (%)** | **1 μM** | **GOF (%)** | **1.5 μM** | **GOF (%)** |
| **Exp 1** | | **replicate 1** | 23.51 | 98.7 | 22.96 | 99 | 25.3 | 97 | 29.36 | 97.8 | 32.18 | 95.4 | 34.88 | 89.8 |
|  |  | **replicate 2** | 23.4 | 98.4 | 22.91 | 98.9 | 25.93 | 98.6 | 30.04 | 96.9 | 32.69 | 93.8 | 33.74 | 88.9 |
|  |  | **replicate 3** | 23.65 | 97.2 | 22.99 | 96.2 | 24.7 | 98.9 | 29.13 | 96.4 | 32.79 | 94.2 | 35.71 | 89.3 |
| **Exp 2** | | **replicate 1** | 22.32 | 90.8 | 22.39 | 93.5 | 23.97 | 99.5 | 29.47 | 96.2 | 31.72 | 93.4 | 34.13 | 93.1 |
|  |  | **replicate 2** | 22.55 | 94.1 | 22.37 | 92.9 | 24.17 | 98.8 | 29.2 | 96.9 | 32.56 | 92.8 | 33.61 | 89.3 |
|  |  | **replicate 3** | 22.7 | 92.6 | 22.32 | 91.7 | 23.93 | 98.8 | 29.33 | 95.4 | 31.6 | 93.5 | 34.47 | 92.9 |

**Table S2. Calculated periods and goodness of fits (GOFs) for oscillations of *Per2:luc* reporter signals in U2OS cells treated with direct circadian modulators and respective controls.**

|  | | **NT** | **GOF (%)** | **DMSO** | **GOF (%)** | **2.7 μM KL001** | **GOF (%)** | **1.5 μM PF-670462** | **GOF (%)** |
| --- | --- | --- | --- | --- | --- | --- | --- | --- | --- |
| **Exp 1** | **replicate 1** | 22.68 | 97.9 | 22.87 | 96.7 | 24.08 | 92.2 | 35.53 | 74.5 |
|  | **replicate 2** | 22.8 | 96.3 | 22.81 | 98 | 28.52 | 93.2 | 32.61 | 86.9 |
|  | **replicate 3** | 22.96 | 96.1 |  |  | 25.16 | 92.9 | 30.87 | 77.5 |
| **Exp 2** | **replicate 1** | 22.89 | 97.5 | 22.7 | 97.8 | 24.53 | 91.8 | 35.1 | 75.4 |
|  | **replicate 2** | 22.77 | 97.7 | 22.66 | 97.6 | 28.64 | 93.3 | 31.23 | 82.3 |
|  | **replciate 3** | 23.53 | 97 |  |  | 2.87 | 93.8 | 30.95 | 83.9 |

**Table S3. Calculated periods and goodness of fits (GOFs) for oscillations of *Bmal1:luc* reporter signals in U2OS cells treated with indirect circadian modulators and respective controls.**

| **SP600125** | | **NT** | **GOF (%)** | **DMSO** | **GOF (%)** | **0.8 μM** | **GOF (%)** | **2 μM** | **GOF (%)** | **7 μM** | **GOF (%)** | **10 μM** | **GOF (%)** |
| --- | --- | --- | --- | --- | --- | --- | --- | --- | --- | --- | --- | --- | --- |
| **Exp 1** | **replicate 1** | 23.03 | 99 | 22.81 | 98.3 | 24.68 | 99.2 | 26.21 | 98.9 | 30.96 | 89.9 | 33.74 | 90.8 |
|  | **replicate 2** | 22.89 | 98.8 | 22.66 | 98.6 | 24.8 | 98.6 | 26.21 | 98.4 | 31.03 | 94.1 | 33.38 | 87.7 |
|  | **replicate 3** | 22.9 | 98.9 | 22.31 | 98.6 | 25 | 98.9 | 26.9 | 98.9 | 30.7 | 93.2 | 34.5 | 92.1 |
| **Exp 2** | **replicate 1** | 23.29 | 97.6 | 23.48 | 96.5 | 24.74 | 97.4 | 26.41 | 97.7 | 31.87 | 87.4 | 34.54 | 83.6 |
|  | **replicate 2** | 23.45 | 97.7 | 23.39 | 96.9 | 24.96 | 98.4 | 26.25 | 97.4 | 31.95 | 86.3 | 33.49 | 85.4 |
|  | **replicate 3** | 23.33 | 98.6 | 23.31 | 96.6 | 24.68 | 97.8 | 26.13 | 98.4 | 31.46 | 87.4 | 33.84 | 85 |
| **Chir99021** | | **NT** | **GOF (%)** | **DMSO** | **GOF (%)** | **0.1 μM** | **GOF (%)** | **0.3 μM** | **GOF (%)** | **0.9 μM** | **GOF (%)** | **3 μM** | **GOF (%)** |
| **Exp 1** | **replicate 1** | 23.19 | 99.3 | 22.8 | 98.2 | 22.6 | 98.8 | 21.72 | 96.4 | 19.79 | 95.1 | 18.35 | 96.5 |
|  | **replicate 2** | 23.2 | 94 | 22.7 | 99.1 | 22.34 | 98.6 | 21.85 | 97 | 19.84 | 96.1 | 18.37 | 83.4 |
|  | **replicate 3** | 23.1 | 98.8 | 22.98 | 98.6 | 22.48 | 98 | 21.8 | 97.9 | 19.66 | 95.5 | 18.74 | 87.2 |
| **Exp 2** | **replicate 1** | 22.82 | 98.9 | 22.8 | 88 | 23.68 | 97.1 | 21.99 | 98.2 | 20.63 | 92.2 | 18.62 | 50.5 |
|  | **replicate 2** | 25.35 | 85.3 | 23.28 | 98.8 | 23.46 | 96 | 22.32 | 96.8 | 20.63 | 92.5 | 18.25 | 74.5 |
|  | **replciate 3** | 24.6 | 97.6 | 22.73 | 97.5 | 22.55 | 98.8 | 22.7 | 96.6 | 20.5 | 90.8 | 18.72 | 94.7 |
| **Etoposide** | | **NT** | **GOF (%)** | **DMSO** | **GOF (%)** | **0.3 μM** | **GOF (%)** | **0.8 μM** | **GOF (%)** | **2 μM** | **GOF (%)** | **7 μM** | **GOF (%)** |
| **Exp 1** | **replicate 1** | 23.19 | 99.3 | 22.8 | 98.2 | 22.95 | 96.6 | 23.82 | 97.4 | 23.38 | 96.9 | 22.25 | 93.1 |
|  | **replicate 2** | 23.2 | 94 | 22.7 | 99.1 | 23.01 | 98.1 | 24.6 | 94.7 | 22.52 | 98.1 | 22.11 | 94.3 |
|  | **replicate 3** | 23.1 | 98.8 | 22.98 | 98.6 | 23.12 | 98 | 23.16 | 98.1 | 22.4 | 98.2 | 22.71 | 94.1 |
| **Exp 2** | **replicate 1** | 22.82 | 98.9 | 22.8 | 88 | 23.52 | 97.8 | 23.18 | 92.5 | 23.4 | 96.2 | 22.14 | 92.3 |
|  | **replicate 2** | 25.35 | 85.3 | 23.28 | 98.8 | 23.14 | 98.1 | 23.11 | 97.2 | 23.22 | 96.9 | 22.2 | 85.5 |
|  | **replciate 3** | 24.6 | 97.6 | 22.73 | 97.5 | 22.82 | 98 | 23.94 | 92.3 | 22.83 | 96.9 | 23.39 | 89.2 |
| **Exp 3** | **replicate 1** | 22.79 | 98.9 | 22.68 | 97.6 | 22.65 | 97.2 | 22.87 | 95.1 | 22.09 | 94.4 | 21.35 | 94 |
|  | **replicate 2** | 23.19 | 98.6 |  |  | 23.77 | 93.4 | 22.75 | 95.1 | 21.57 | 94.3 | 21.58 | 93.2 |
|  | **replciate 3** | 22.93 | 98.3 |  |  | 22.52 | 96.8 | 23.79 | 93.9 | 22.02 | 94.4 | 21.18 | 94.4 |

**Table S4. Calculated periods and goodness of fits (GOFs) for oscillations of *Per2:luc* reporter signals in U2OS cells treated with indirect circadian modulators and respective controls.**

|  | | NT | GOF (%) | DMSO | GOF (%) | 10 μM SP600125 | GOF (%) | 3 μM Chir99021 | GOF (%) | 7 μM Etoposide | GOF (%) |
| --- | --- | --- | --- | --- | --- | --- | --- | --- | --- | --- | --- |
| **Exp 1** | **replicate 1** | 23.47 | 98.8 | 22.99 | 97.7 | 32.3 | 84.5 | 18.76 | 76.3 | 21.94 | 97.8 |
|  | **replicate 2** | 23.44 | 98.6 | 22.96 | 98.2 | 31.28 | 84.3 | 18.78 | 93.9 | 21.99 | 97.9 |
|  | **replicate 3** | 23.67 | 97.5 |  |  | 34.07 | 83.2 | 18.6 | 82.8 | 21.96 | 7.7 |
| **Exp 2** | **replicate 1** | 23.52 | 97.6 | 23.12 | 98.8 | 30.69 | 88.1 | 18.81 | 81.1 | 22.08 | 97.5 |
|  | **replicate 2** | 23.4 | 98.9 | 23.17 | 98.4 | 31.72 | 85.1 | 18.96 | 78.1 | 22.22 | 97 |
|  | **replicate 3** | 25.37 | 96.9 |  |  | 31.81 | 85.9 | 19.16 | 91.6 | 22.03 | 98.1 |
